# Plaque Size Tool: an automated plaque analysis tool for simplifying and standardising bacteriophage plaque morphology measurements

**DOI:** 10.1101/2021.04.12.439404

**Authors:** Ellina Trofimova, Paul R Jaschke

## Abstract

Bacteriophage plaque size measurement is essential for phage characterisation, but their manual size estimation requires a considerable amount of time and effort. In order to ease the work of phage researchers, we have developed an automated command-line application called Plaque Size Tool (PST) that can detect plaques of different morphology on the images of Petri dishes and measure plaque area and diameter. Plaque size measurements using PST showed no difference to those obtained with manual plaque size measurement in Fiji, indicating future results using PST are backwards compatible with prior measurements in the literature. PST can be applied to a range of lytic bacteriophages producing oval-shaped plaques, including bull’s-eye morphology. The application can also be used for titer calculation if most of the plaques are stand-alone. As laboratory automation becomes more commonplace, standardised and flexible open-source analytical tools like PST will be important parts of biofoundry and cloud lab bacteriophage workflows.

## INTRODUCTION

Bacterial viruses, or bacteriophages, when co-plated with susceptible hosts produce clearings, or plaques, of different morphology on the lawn of bacterial cells. Each plaque usually represents an infection by one virus particle (1). The common plaque types include clear plaques, plaques with clear centres and turbid edges (bull’s-eye), plaques with a halo, and turbid plaques (2).

Plaques are usually visualised by co-plating phage with appropriate bacterial host strains within solid media using techniques that limit phage diffusion such as using a layer of molten agar or agarose matrix (overlay) (3). After incubation at a particular temperature, clearings on the bacterial lawn become visible in the overlay and available for assessment and phage isolation. Traditionally, plaque size was measured with a ruler or caliper (4) where the accuracy and precision of the measurement depends on the skill and experience of the operator.

Besides being an irreplaceable part of phage isolation procedures plaque morphology is essential for phage phenotypic measurements, including such fitness traits as burst size, adsorption rate, and lysis time (2), and the rate of phage diffusion within a particular medium (5). It is also possible to make a conclusion on the presence or absence of mutations in the phage genome (6) based on changes in plaque morphology, including in rationally engineered phages (7–10).

Currently, there are several methods for semi-automated measurement of bacteriophage plaque sizes using digital images of Petri dishes. Adobe Photoshop’s Ruler Tool is distributed with the default software configuration and allows sizing plaque diameters in pixels but requires a manual measurement. Two additional options ‘Record Measurements’ and ‘Count tool’ might ease the selection of multiple objects but are only available in the extended version of Adobe Photoshop, and also require manual selection (8). Alternatively, there are several free applications available for viral plaque size measurement, but they all have limitations when processing bacteriophage plaques.

One of the free phage plaque size measurement tools is Fiji (11). With ‘Oval’ and ‘Line’ selection tools, it is possible to manually measure the area and diameter of a plaque. The ViralPlaque (12) Fiji plugin is designed to automatically detect plaques, but is only able to detect plaques with clear morphology and cannot process phages that produce plaques with clear centers and turbid edges (bull’s-eye morphology) like coliphage φX174 and *Acinetobacter Baumannii* phage IME200. For clear plaques, to achieve better plaque recognition by the software the user is also required to adjust images manually using several settings like Brightness/Contrast, Blur, and Thresholding (12).

Additionally, several automated and semi-automated applications were created to obtain statistics on mammalian viral plaques, such as Viridot (13), Plaque 2.0 (14), Infection Counter (15). However, these tools are generally used to measure the *number* of viral plaques, not their *morphology*. Furthermore, some of these tools require input images obtained using specific conditions (e.g. fluorescence microscopy) and are not suitable for more general uses like measuring bacteriophage plaques from Petri dishes. Thus, there is currently a lack of tools capable of automatically detecting and measuring bacteriophage plaques to assess their morphology. To solve this issue, we created *Plaque Size Tool* (*PST*) that is able to detect non-overlapping bacteriophage plaques on Petri dish images and measure their characteristics including area and diameter, in a fully automated mode.

## METHODS

### Bacteriophage Plaque Formation

A range of wild-type and mutant φX174 bacteriophage were plated using the double agar overlay method as previously described (10) with *E. coli* C122 (Public Health England NCTC122) grown in Phage LB (16) on 90 mm Petri dishes. The plates were incubated overnight at 37° C.

### Bacteriophage plate imaging

Images of plates containing plaques of different size (one plate per one image) were taken using Bio-Rad Gel Doc XR+ Gel Documentation system and Image Lab software version 6.0.1 with a standard filter, white epi-illumination options and the image exposure time equal to 0.750 sec. Images were exported in TIFF format in 300 (10 images) and 600 (7 images) dots per inch (DPI) resolution. The dimensions of 300 DPI plates were 1606×1200 pixels, 600 DPI plates were 2811×2100 pixels.

### Manual plaque size measurements

Plaques were manually identified and measured with Fiji version 2.1.0/1.53c using the ‘Analyze/Measure/Oval’ feature. Each image was magnified up to 400% prior to the measurement, and plaques were manually circled with the ‘Oval’ feature. The identifiers of plaques were assigned in the same order as on the PST output image using ‘Analyze/Tools/ROI Manager’. After all plaques were selected, their ‘Area’ measurements were taken using the Region of Interest (ROI) Manager and exported into CSV for further analysis.

### Statistical analysis

To compare automatically detected plaque area values using PST with manually measured plaques in Fiji, plate images were divided into three groups: 300 DPI (n=10), 600 DPI (n=7), and all plates (n=17). Obtained plaque size values were tested for normal data distribution with the Shapiro-Wilk normality test in ‘jamovi’ (17). Plaque area values in square pixels were compared between PST and Fiji in pairs where the same plaque had the same identifier in PST and Fiji. The Wilcoxon signed rank test was used to compare the groups with Python 3 (18) scripting using the module ‘wilcoxon’ from the package scipy.stats (version 1.5.4) (19). For small plaque size determination, an average diameter of plaques was calculated from the plates that contained noticeably smaller plaques and were measured with the ‘-small’ flag.

The number of correctly identified plaques by Plaque Size Tool was measured for each plate to compare with the total number of non-overlapping plaques counted manually and obtain the tool accuracy. All extra objects detected as plaques (false positives), along with incorrectly measured plaques, were considered as not accurately identified and not included in the comparison of area values. PST accuracy values for the average PST plaque diameter calculated for each plate were analysed using a linear regression method (linregress) from the package scipy.stats (version 1.5.4) (19).

### Plaque Size Tool development and distribution

The command-line tool PST was written in Python 3 using the Open Source Computer Vision Library (OpenCV) ver. 4.5.1.48 library for object detection (20). The tool is available on GitHub at https://github.com/ellinium/plaque_size_tool under Apache License 2.0. The installation documentation is specified in the manual (File S1) and on the GitHub webpage.

PST currently receives three input arguments (also called flags): (1) the full path to an image (-i) or full path to a directory with images (-d) (required), (2) plate size in millimetres (-p) (optional), and (3) a flag for processing small plaques (-small) (optional). The tool supports images containing one Petri dish in JPG, JPEG, TIF, TIFF and PNG formats.

### Plaque Size tool image processing workflow

All following methods of image transformation belong to OpenCV–Python library (21). First, each image that contains only one Petri dish is converted to grayscale mode and denoised using the non-local means denoising algorithm (Fig. 1). After that, the image is blurred using a Gaussian blur method. The output image is then converted to an 8-bit image, and its contrast is increased. In order to detect the contours of the plaques, adaptive threshold and Laplacian transformation are applied. A binary image with oval contours is then formed. The image is processed further to find all image contours in a two-level hierarchy structure that have all external contours (i.e. its boundary) placed in hierarchy 1 and all the contours inside an object in hierarchy 2.

**Fig. 1.**
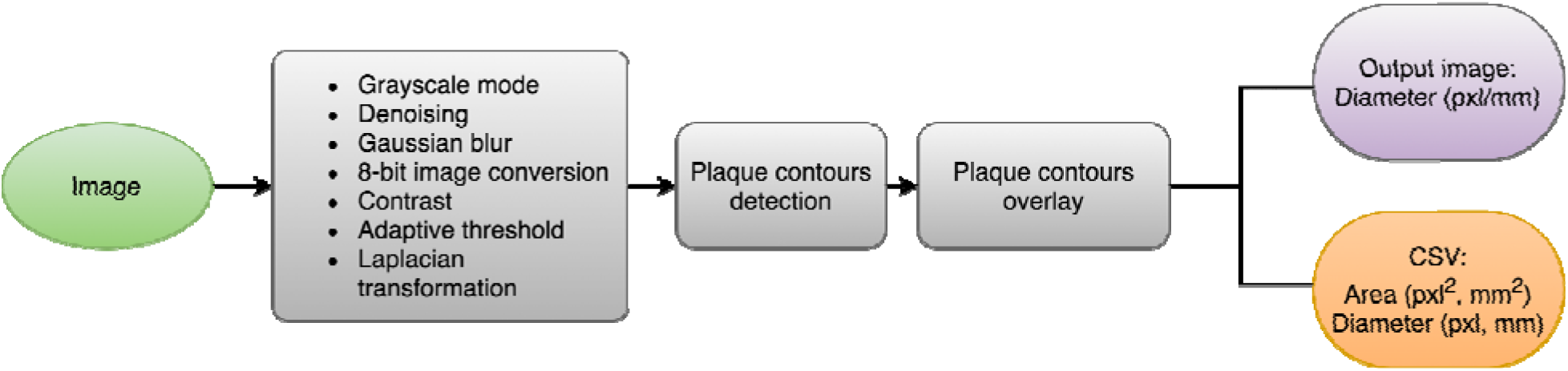
Workflow of Plaque Size Tool. The green filling colour indicates input files, grey indicates image-processing steps. Purple and orange colours indicate output files. Pxl (pixel), mm (millimetre).

After all contours are found, convex hulls are drawn for a set of points of each contour. Based on the hull values, a plaque area is calculated in square pixels (pxl^2^). A plaque diameter is also calculated in pixels using the formula 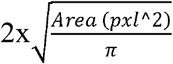. The Petri dish contour is used to convert the area and diameter of plaques from pixels to millimetres if the ‘-p’ (plate size) flag is specified.

### Plaque Size Tool output files

The detected plaque contours are drawn on the high-contrast image in green colour and saved into an output folder ‘out’ that is created automatically in the directory PST is launched from. Additional information is saved into a comma-separated file (CSV) and contains plaque identifiers (INDEX_COL) that corresponds to the identifier of the same plaque on the output image, plaque area in pixels (AREA_PXL), and plaque diameter in pixels (DIAMETER_PXL). If the −p flag is used during PST execution, then plaque area in millimetres (AREA_MM), and plaque diameter in millimetres (DIAMETER_MM) are also returned in the CSV file.

## RESULTS

### Plaque Size Tool accuracy and detection capability

Plaque Size Tool (PST) was designed to measure non-overlapping plaques from digital images of double agar overlay plates. The command-line tool was developed in Python 3 using Open Source Computer Vision Library (OpenCV). The tool is run with three input arguments: (1) the full path to an image (-i) or full path to a directory with images (-d) (required), (2) plate size in millimetres (-p) (optional), and (3) a flag for processing small plaques (-small) (optional).

Although PST is able to identify the contours of overlapping plaques, we chose to not use these plaques for determination of plaque morphology because the size of a plaque can be affected by its adjacent neighbour. The output of the tool is an image with recognized and measured plaques circled and labelled with a hash (#) symbol, unique identifier separated from diameter value with a full colon, and a CSV file containing area and diameter values for every identified plaque (Fig. 2).

**Fig. 2.**
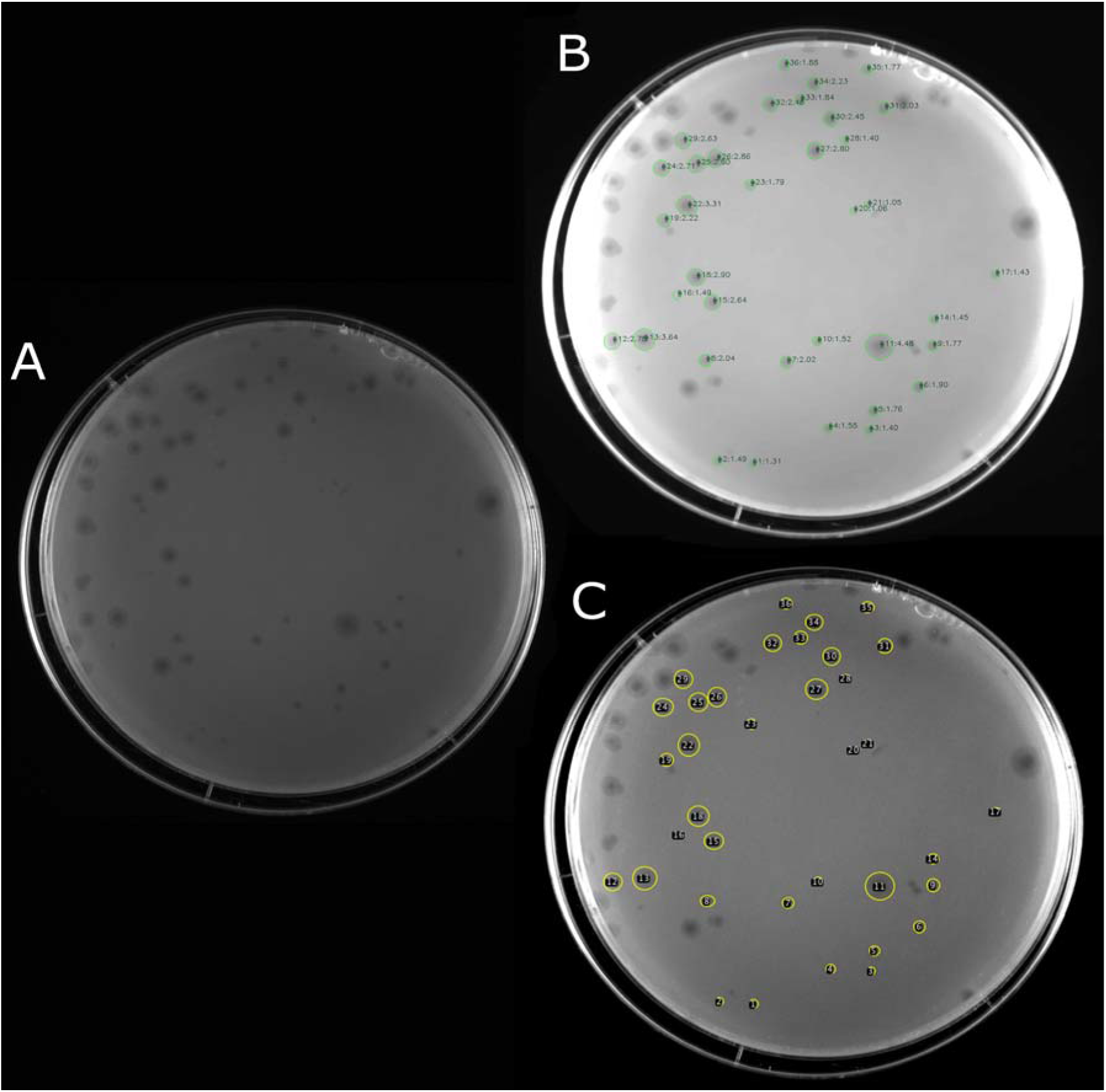
Petri plates with bacteriophage plaques before and after analysis with plaque measurement tools. (A) The original image in TIFF format. (B) An output image created by Plaque Size Tool. Detected plaques are circled with green colour and labelled with a hash symbol (#) followed by a unique numeric identifier and the plaque diameter in mm (e.g. #23:1.79). (C) Manual plaque selection in Fiji of the same image.

PST was first tested by processing 17 Petri dish images in 300 and 600 DPI resolution TIFF format from experiments on phage φX174 plaques in our lab (Table S1). Seven plates containing visibly small plaques were processed with the ‘-small’ flag. The images contained a total of 601 non-overlapping plaques as identified by manual measurements. On average PST was able to identify and measure 83.0% (499/601) of plaques across the dataset (Table S3). At the plate level, detection accuracy varied between 52-100% (Table S1).

Detection accuracy did not correlate with plaque size. Mean PST plaque diameter was measured for each plate and this was compared to the PST detection rate. Using linear regression analysis against these data we found no correlation between the PST plaque detection rate and the average plaque diameter on a Petri dish (R^2^=0. 02, slope=1.18) (Fig. 3).

**Fig. 3.**
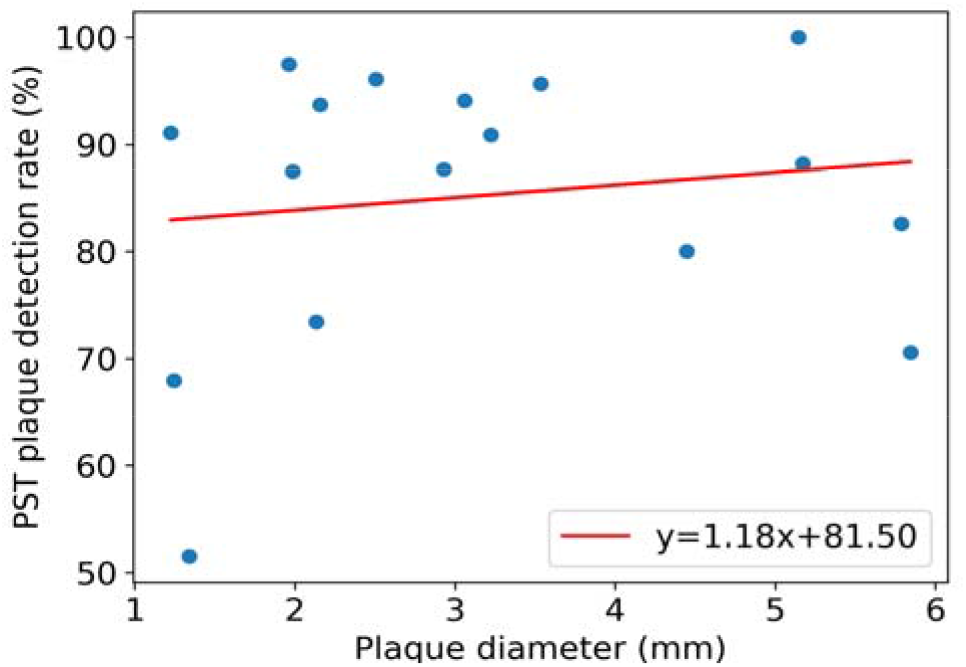
Correlation between PST plaque detection rate and the average plaque diameter. The linear least-squared fit equation is provided for the method.

### Comparison to manual plaque measurement method

We next benchmarked the PST against Fiji to determine how similar the calculated morphology values between these two tools are. The values of area obtained by PST from the CSV output file for each Petri dish were compared with the area values obtained by manual plaque measurement on the same Petri dish with ‘Analyze/Measure/Oval’ feature in Fiji. Comparing the distribution of measured plaque sizes between PST and Fiji showed the two methods generate very similar results (Fig. 4).

**Fig. 4.**
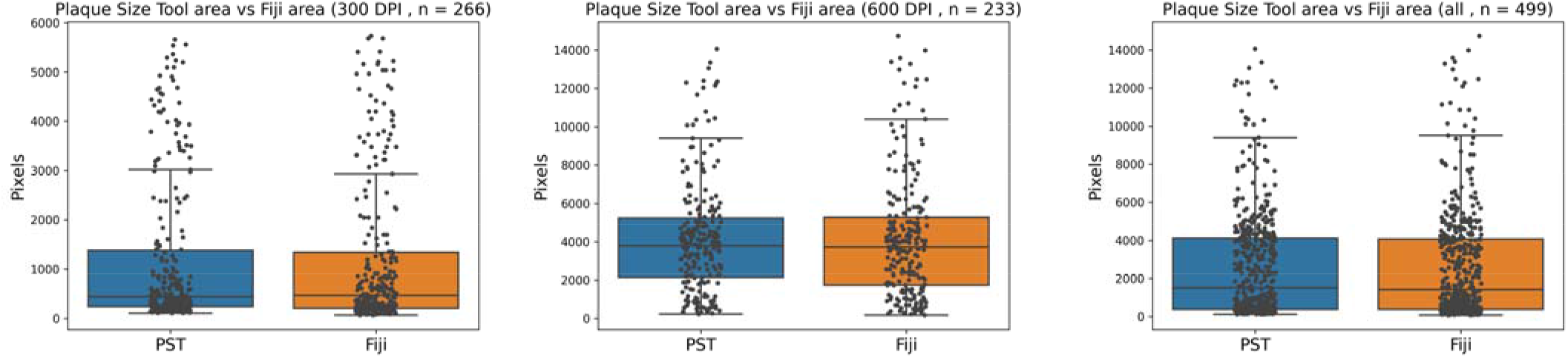
Measured plaque areas from automated PST and manual Fiji analysis shows no statistical difference. (A) 300 DPI. (B) 600 DPI. (C) All. Box plots for the three plate groups with individual data points overlaid. Significance of plaque area difference between conditions was measured using the Wilcoxon signed rank test (18).

As the plaque area measurements are not normally distributed (Table 1) we used the Wilcoxon signed rank test to compare the sizes of non-overlapping plaques measured automatically by PST paired with the manually measured in Fiji. The comparison showed there was no significant difference between the plaque sizes measured by PST and Fiji (Fig. 4) at the 0.05 significance level (Table 2).

**Table 1.** Tests for normality of PST plaque area values.

| Image Group | W value | p-value |
| --- | --- | --- |
| 300 DPI | 0.926 | <0.001 |
| 600 DPI | 0.962 | <0.001 |
| All | 0.968 | <0.001 |

The normality of data distribution was tested with the Shapiro-Wilk normality test in 'jamovi' (17).

**Table 2.** Wilcoxon signed rank test results for PST compared to manual Fiji plaque area measurements.

| Image Group | W value | p-value |
| --- | --- | --- |
| 300 DPI | 24515.500 | 0.674 |
| 600 DPI | 13397.000 | 0.821 |
| All | 98774.500 | 0.283 |

The Wilcoxon signed rank test was used to compare the groups with Python 3 (18) scripting using the module 'wilcoxon' from the package scipy.stats (version 1.5.4) (19).

### The effect of image resolution and type on plaque detection

To determine the effect of image resolution on plaque detection rate, seven plates that were previously analysed from 600 DPI resolution TIFF images group were instead exported as 300 DPI TIFF images and analysed with PST. We found on average that the number of detected plaques in the 300 DPI images was reduced from the 600 DPI images by 31 %. The reduction ranged from 4.4 % to 57.7%, depending on the plate, without correlation with the plaque size (R^2^=0.0001, slope=0.17).

To test the ability of PST to measure plaque sizes from a range of digital images and phage we downloaded seven random plate images from the Actinobacteriophage database (22). The images were captured by a range of different cameras, including mobile phones, and had a range of different resolutions (Table S2). For the seven additional Actinobacteriophage plates, PST was able to detect 88.1% of plaques on the plates with plaques larger than 1 mm (133/151). On plates of Arthrobacter phage Rizwana plaques with sizes less than 1 mm PST detected 54.5 % (12/22) using the -small flag. A high proportion of plaques, 72-88%, were detected on the JPG and PNG plate images of low resolution and size (150 DPI and 413×310 pixels, 72 DPI and 624×462 pixels, and 144 DPI and 646×626 pixels) indicating that PST application can effectively process images of low resolution and size from a range of devices (Table S2).

## DISCUSSION

Manual measurement of phage plaque dimensions can be tedious, time-consuming, lead to erroneous calculations, and is non-standardised. In order to ease the work of phage scientists, we created an automated command-line tool for phage plaque detection and assessment of their diameter and area parameters. Easily installed Plaque Size Tool, written in Python 3, allows single and batch image processing of bacteriophage plaques on Petri dishes in a wide range of resolutions and file formats. The tool can be applied to various lytic phages producing plaques of different morphology: clear and turbid plaques, plaques with halo, and plaques with clear centres and turbid edges (bull’s-eye morphology). Only non-overlapping plaques are included in the measurement as the size of merged plaques is affected by their neighbour.

The tool can detect between 70 and 100 % of plaques on a Petri dish with plaques of diameter >1.5 mm, and 51-91 % of plaques <1.5 mm. The difficulty for plaque detection occurs on the edges of Petri dishes due to the change of an image exposure during its processing. However, even for plaques less than 1 mm, the detected plaque number on most Petri dish images is sufficient for robust calculation of size, especially considering that such plaques are very difficult to measure manually. For TIFF images, resolution lower than 600 DPI decreases plaque detection rates by PST but other image formats seem to perform better than TIFF at a range of lower resolutions which might be linked to the original contrast and brightness of the image. If an image simultaneously has a low resolution and size, the ‘-small’ flag can be used to process such image with PST.

PST is highly accurate if compared with measurements performed manually, and the results are reproducible for each processed image as the detection algorithm will always produce the same plaque count and size measurements from the same input image.

Plaque size measurements using PST were statistically indistinguishable from those we measured with manual plaque size measurement tool Fiji (Fig. 4). As a result, PST measurements can be directly compared with past manual Fiji measurements, enabling backwards compatibility with prior literature values.

The tool can also be used for phage titer calculation with no modification if most of the plaques on the plate are not overlapping or around the edge. Alternatively, a user can manually calculate a number of plaques that were not labelled by PST (i.e., merged plaques or those around the edge) and add it to the total number of plaques calculated by the tool.

To summarise, we have created a tool that addresses a gap within the current offerings for automated phage plaque analysis. It requires minimal user actions and can process a range of Petri dish images of different format and resolution in single and batch mode. Computational tools such as PST will be an increasingly important component within analytical pipelines for phage engineering and novel phage isolation and characterisation in cloud labs and biofoundries in the future (23,24).

## Acknowledgements

We thank Ilya Trofimov for elucidating features of OpenCV library and Russell M Vincent for helpful discussions. Russel M Vincent and Bradley W Wright provided the φX174 bacteriophage images.

## Authors and contributors

The contributions of authors of this work according to the CRediT contribution taxonomy were: ET: conceptualization; data curation; formal analysis; investigation; methodology; software; visualization; writing-original draft; writing-review and editing.

PRJ: conceptualization; funding acquisition; project administration; supervision; writing-review and editing.

