## Supplementary material for "Plaque Size Tool: an automated plaque analysis tool for simplifying and standardising bacteriophage plaque morphology measurements": -

Affiliations:
[1] Department of Molecular Sciences, Macquarie University, Balaclava Rd, Macquarie Park NSW 2109, Australia

**CONTENTS**

SUPPORTING TABLES

Table S1: ΦX174 Petri dish image characteristics

Table S2: Image characteristics of additional phage plates from Actinobacteriophage database

Table S3: Original and Plaque Size Tool processed ΦX174 Petri dish images

SUPPORTING FILES

File S1: Plaque Size Tool User Manual

**Table S1. ΦX174 Petri dish image characteristics.**

| Image name | PST flag | Resolution (DPI) | Image size (pixels × pixels) | Average Plaque Size (mm) | Plaque number (manual count) | Plaque number (PST) | False positive plaques | Incorrect plaques | PST correct plaques (overall) | PST accuracy (%) |
|---|---|---|---|---|---|---|---|---|---|---|
| Plate_1.tif | small | 600 | 2811 x 2100 | 1.22±0.27 | 45 | 44 | 3 | 0 | 41 | 91.11 |
| Plate_2.tif | none | 600 | 2811 x 2100 | 5.17±0.38 | 17 | 19 | 0 | 4 | 15 | 88.24 |
| Plate_3.tif | none | 600 | 2811 x 2100 | 3.06±0.60 | 51 | 51 | 2 | 1 | 48 | 94.12 |
| Plate_4.tif | none | 600 | 2811 x 2100 | 3.22±0.36 | 11 | 10 | 0 | 0 | 10 | 90.91 |
| Plate_5.tif | none | 600 | 2811 x 2100 | 2.93±0.45 | 57 | 51 | 1 | 0 | 50 | 87.72 |
| Plate_6.tif | none | 600 | 2811 x 2100 | 3.53±1.01 | 47 | 48 | 0 | 3 | 45 | 95.74 |
| Plate_7.tif | none | 600 | 2811 x 2100 | 2.51±0.35 | 26 | 25 | 0 | 0 | 25 | 96.15 |
| Plate_8.tif | none | 300 | 1606 x 1200 | 5.14±1.20 | 19 | 19 | 0 | 0 | 19 | 100.00 |
| Plate_9.tif | small | 300 | 1606 x 1200 | 1.96±0.57 | 41 | 45 | 5 | 0 | 40 | 97.56 |
| Plate_10.tif | none | 300 | 1606 x 1200 | 5.79±0.90 | 23 | 25 | 1 | 5 | 19 | 82.61 |
| Plate_11.tif | small | 300 | 1606 x 1200 | 1.25±0.29 | 53 | 38 | 1 | 2 | 35 | 66.04 |
| Plate_12.tif | none | 300 | 1606 x 1200 | 4.45±0.61 | 15 | 12 | 0 | 0 | 12 | 80.00 |
| Plate_13.tif | small | 300 | 1606 x 1200 | 1.34±0.33 | 66 | 37 | 3 | 0 | 34 | 51.52 |
| Plate_14.tif | none | 300 | 1606 x 1200 | 5.85±0.51 | 17 | 17 | 0 | 5 | 12 | 70.59 |
| Plate_15.tif | small | 300 | 1606 x 1200 | 2.14±0.76 | 49 | 36 | 0 | 0 | 36 | 73.47 |
| Plate_16.tif | small | 300 | 1606 x 1200 | 2.16±0.72 | 48 | 46 | 1 | 0 | 45 | 93.75 |
| Plate_17.tif | small | 300 | 1606 x 1200 | 1.99±0.91 | 16 | 17 | 3 | 0 | 14 | 87.50 |

Columns: 'False positive plaques' - additional incorrect objects that were detected on a plate as plaques (bubbles, dark spots and etc.). 'Incorrect plaques' - plaques that were measured inaccurately (merged plaques, a partial selection of a plaque, larger/smaller plaque size). 'PST correct plaques (overall)' - The overall plaque number calculated by PST ('Plaque number (PST)) minus 'False positive plaques' and 'Incorrect plaques'.

**Table S2. Image characteristics of additional phage plates from Actinobacteriophage database.**

| Phage name | File URL | Dimensions | DPI | File Format | Image colour | Device | PST flags | Plaque number (manual count) | Plaque number (PST) | Correctly detected plaques (%) |
|---|---|---|---|---|---|---|---|---|---|---|
| Mycobacterium phage DS6A | https://phagesdb.org/media/plaquePics/2001-07-23_DS6A_on_BCG.jpg | 624 × 462 | 72 | JPG | greyscale | unknown | small | 25 | 23 | 92.00 |
| Streptomyces phage Aaronocolus | https://phagesdb.org/media/plaquePics/Aaronocolus_Plaque_2.jpg | 1360 × 1024 | 72 | JPG | greyscale | unknown | small | 37 | 31 | 83.78 |
| Mycobacterium phage Durfee | https://phagesdb.org/media/plaquePics/Durfee_Plaque.png | 413 × 310 | 150 | PNG | greyscale | unknown | small | 18 | 13 | 72.22 |
| Arthrobacter phage Nitro | https://phagesdb.org/media/plaquePics/Nitro_Plaque.JPG | 3024 × 4032 | 72 | JPEG | colour | mobile phone camera | none | 10 | 10 | 100.00 |
| Arthrobacter phage Rizwana | https://phagesdb.org/media/plaquePics/Rizwana_Plaque.jpg | 2814 × 2825 | 72 | JPG | colour | mobile phone camera | small | 22 | 12 | 54.55 |
| Mycobacterium phage Thresher | https://phagesdb.org/media/plaquePics/Thresher_Plaque.jpg | 1836 × 3264 | 72 | JPG | colour | mobile phone camera | none | 13 | 12 | 92.31 |
| Streptomyces phage Katalie | https://phagesdb.org/media/plaquePics/Katalie_Plaque.png | 646 × 626 | 144 | PNG | colour | unknown | small | 25 | 22 | 88.00 |
| Gordonia phage Anaysia | https://phagesdb.org/media/plaquePics/Anaysia_Plaque_dMih9qg.jpg | 3024 × 3024 | 72 | JPG | colour | mobile phone camera | none | 23 | 22 | 95.65 |

**Table S3. Original and Plaque Size Tool processed ΦX174 Petri dish images**

| Plate name | Original image (TIFF) | Plaque Size Tool output image |
|---|---|---|
| Plate_1.tif | 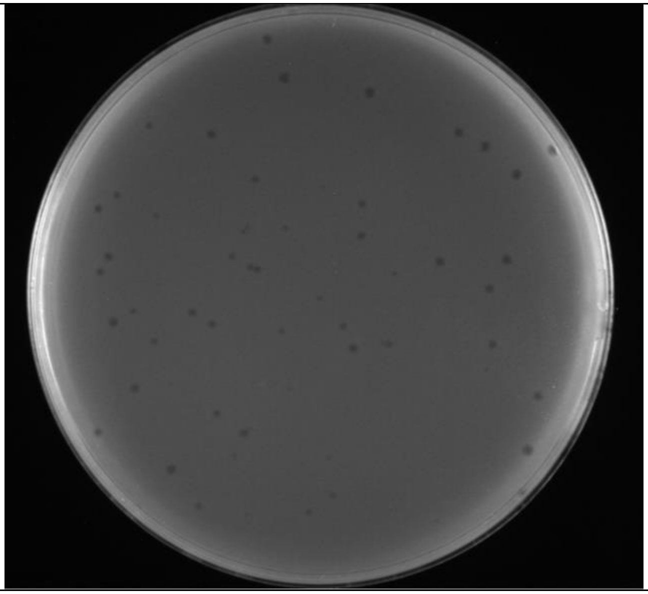 | 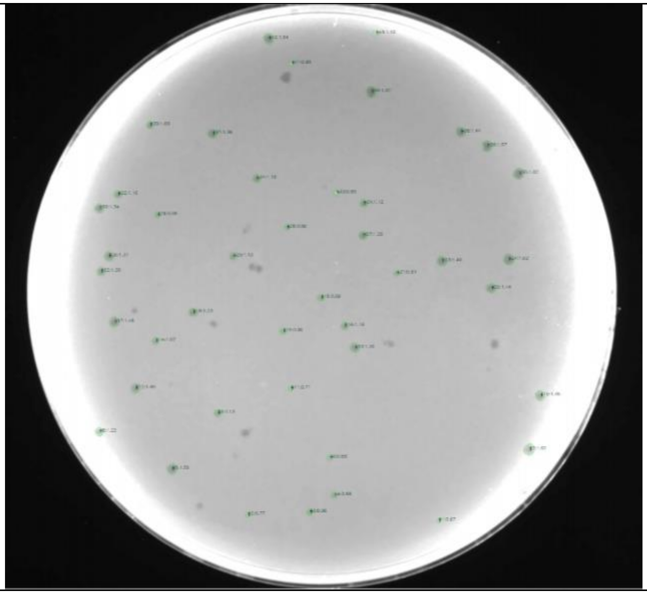 |

| | | |
|---|---|---|
| **Plate_2.tif** | 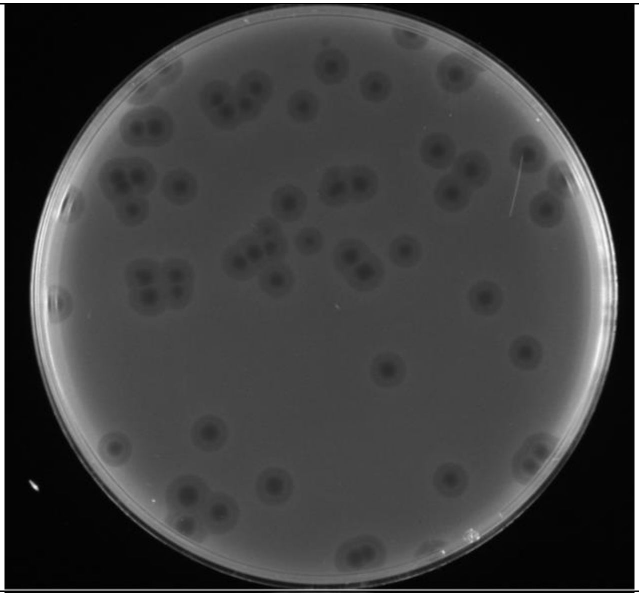 | 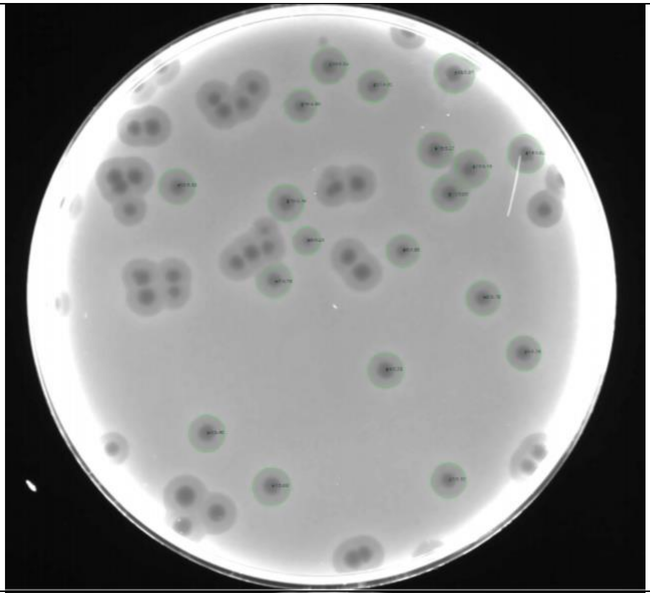 |
| **Plate_3.tif** | 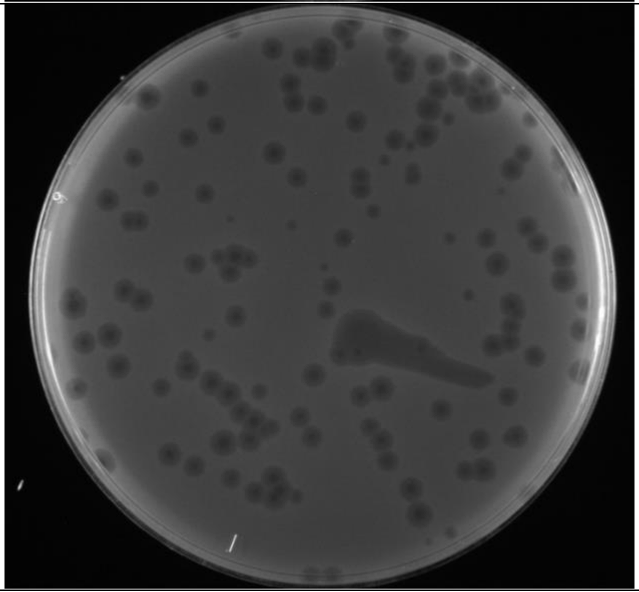 | 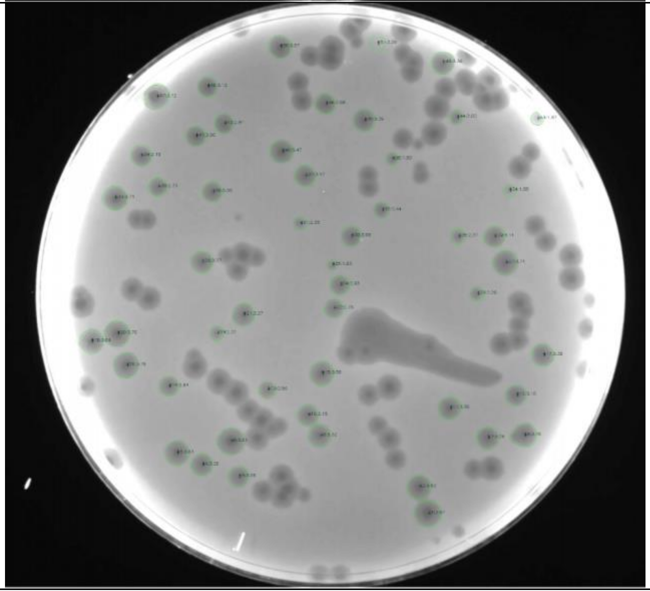 |

| | | |
|---|---|---|
| **Plate_4.tif** | 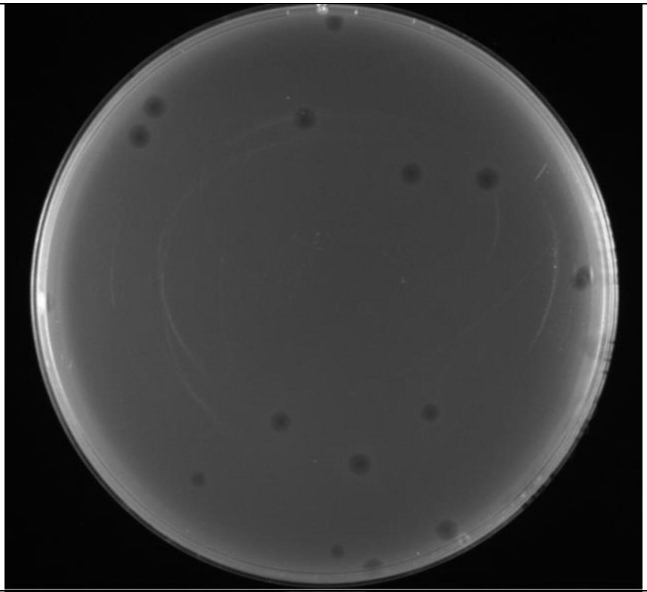 | 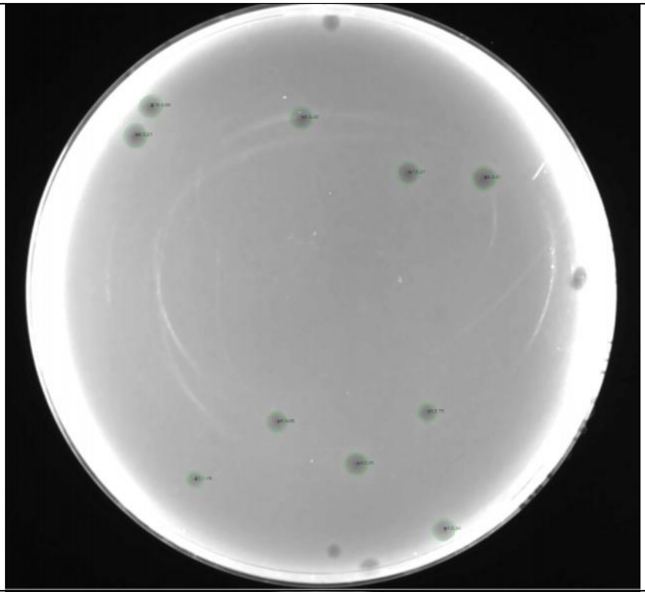 |
| **Plate_5.tif** | 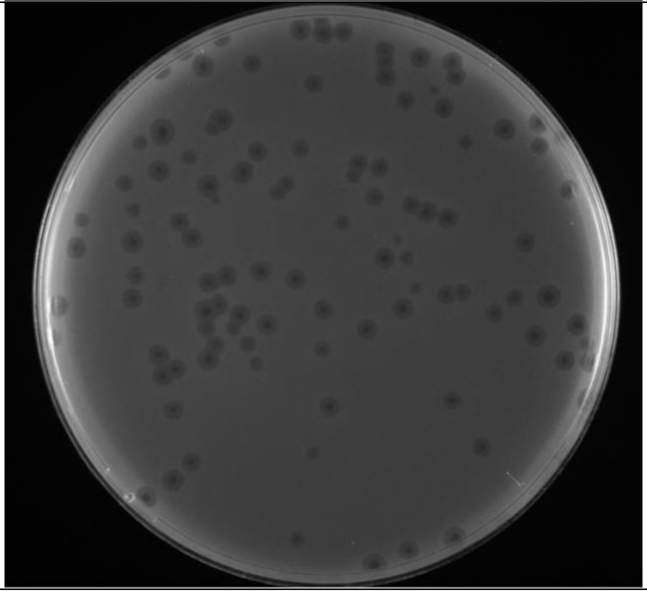 | 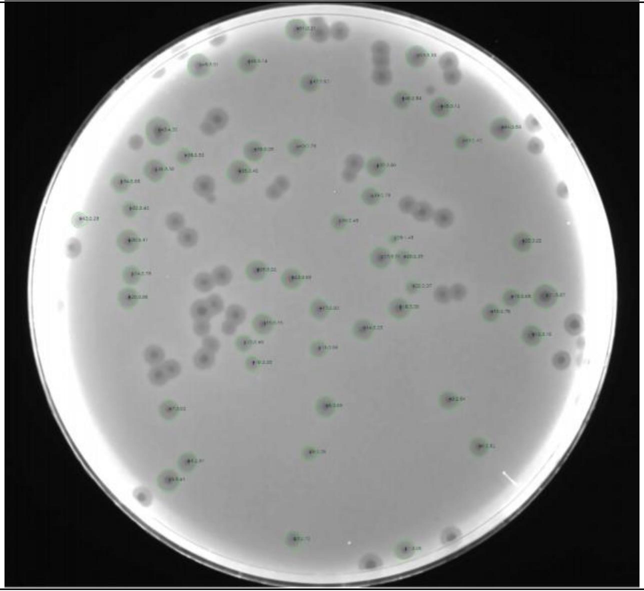 |

| | | |
|---|---|---|
| **Plate_6.tif** | 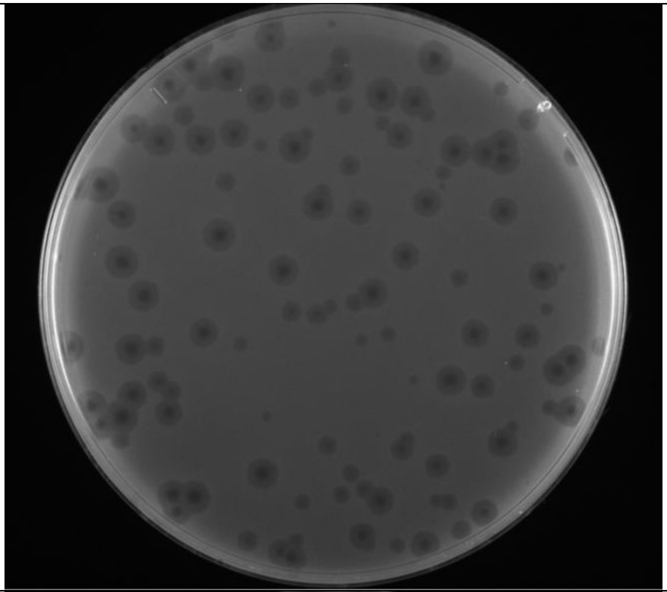 | 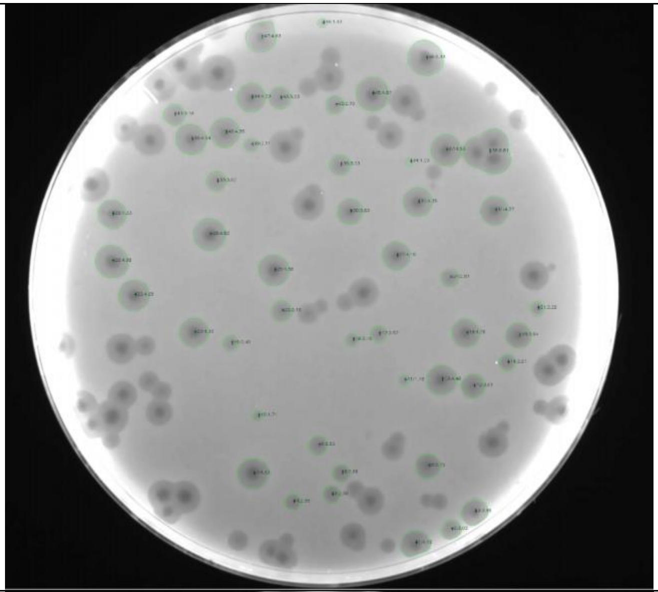 |
| **Plate_7.tif** | 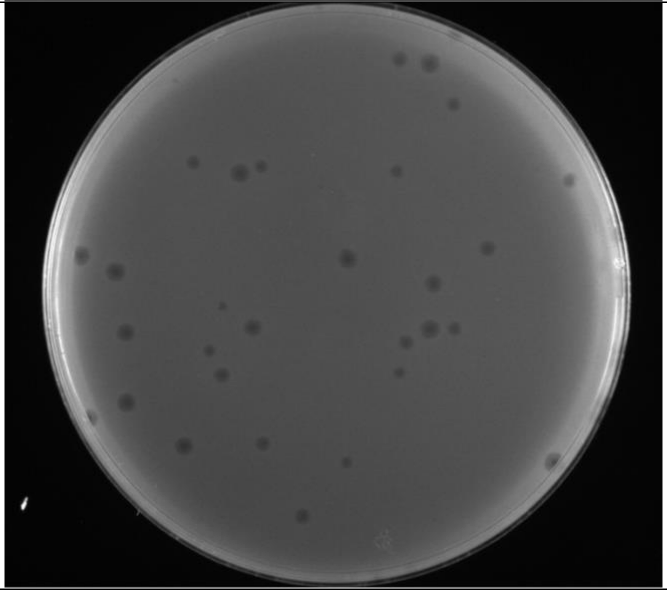 | 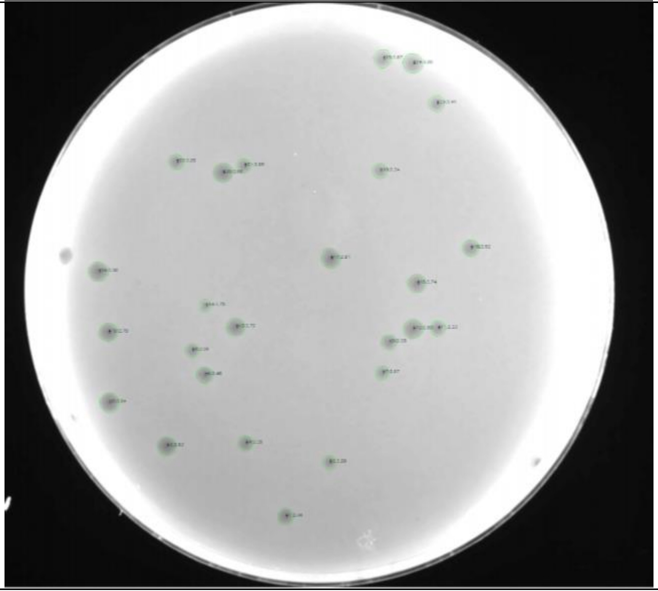 |

| | | |
|---|---|---|
| **Plate_8.tif** | 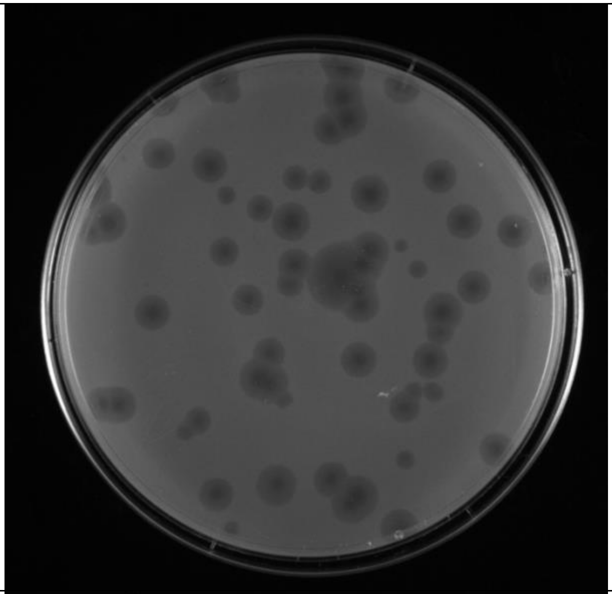 | 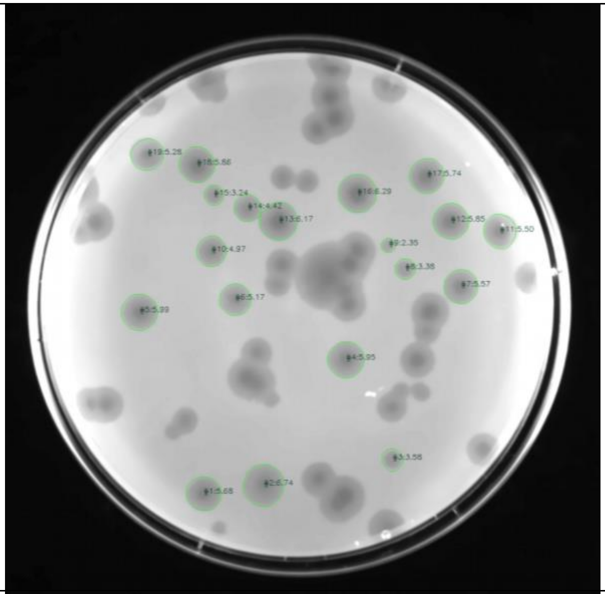 |
| **Plate_9.tif** | 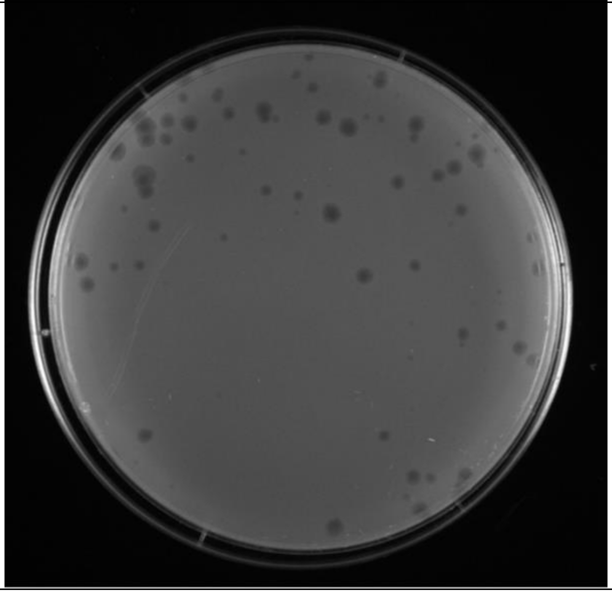 | 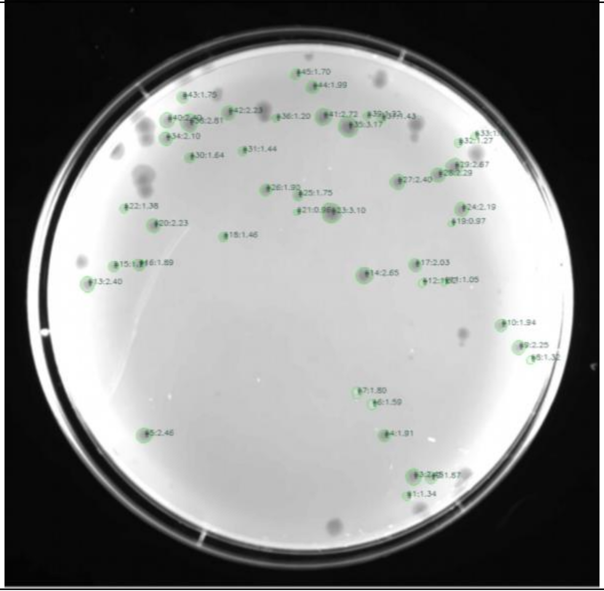 |

| | | |
|---|---|---|
| **Plate_10.tif** | 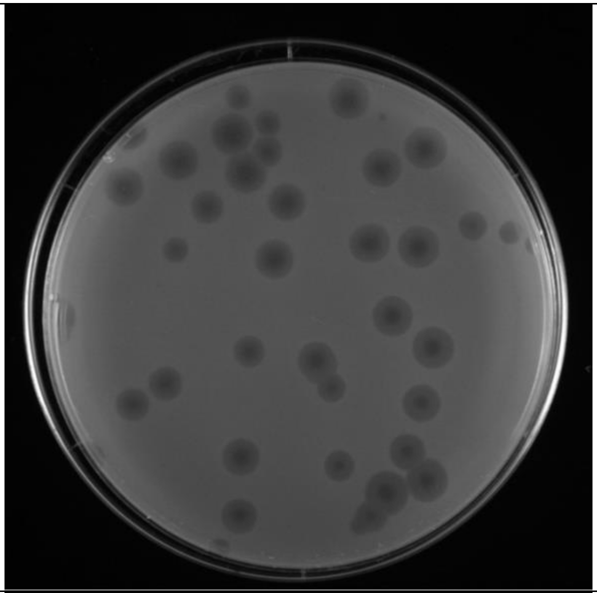 | 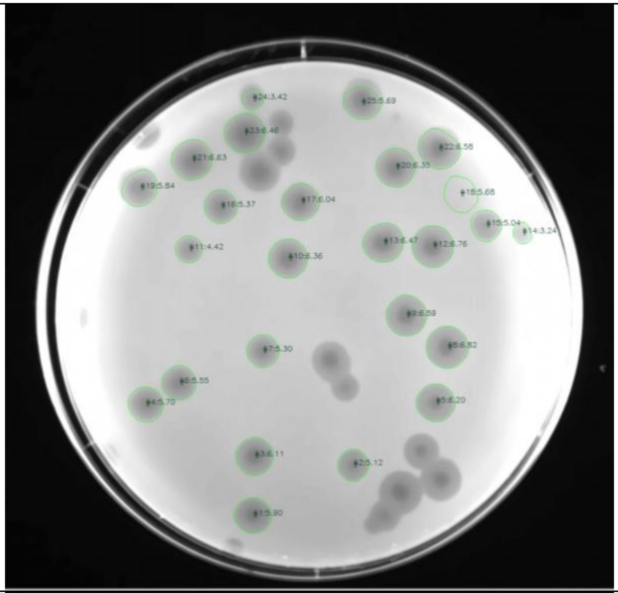 |
| **Plate_11.tif** | 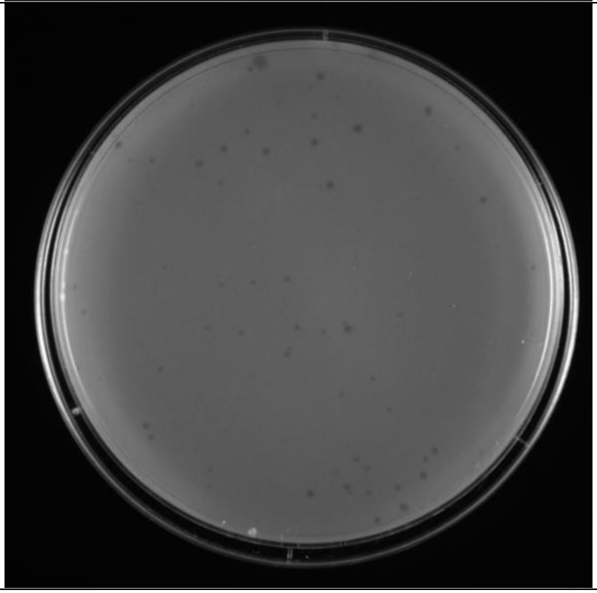 | 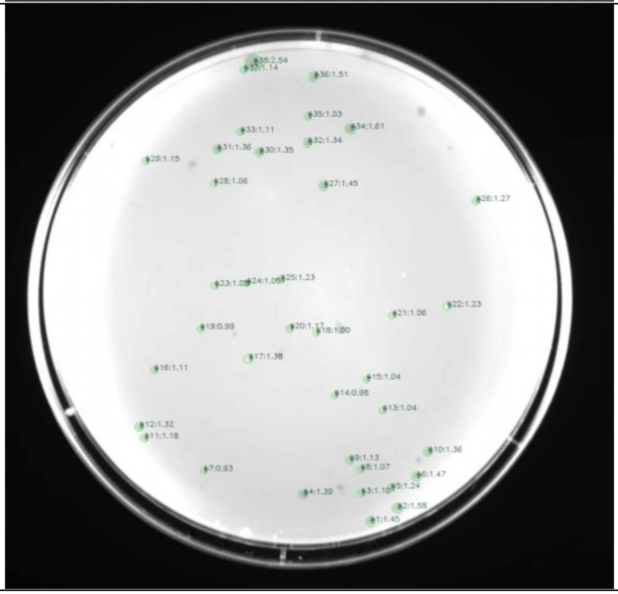 |

| | | |
|---|---|---|
| **Plate_12.tif** | 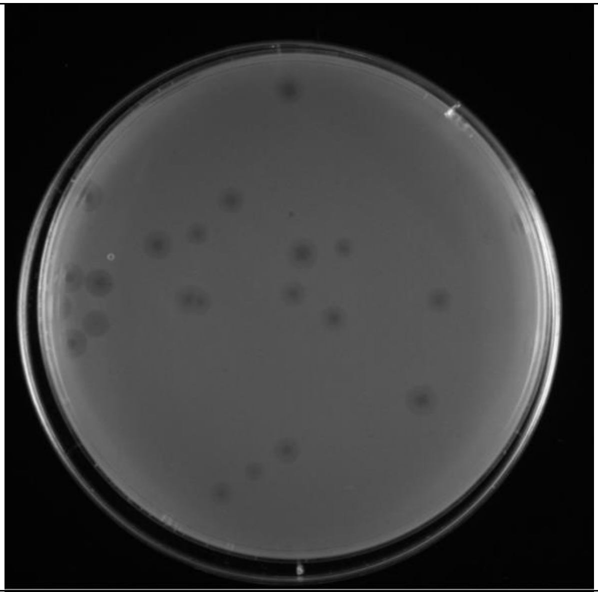 | 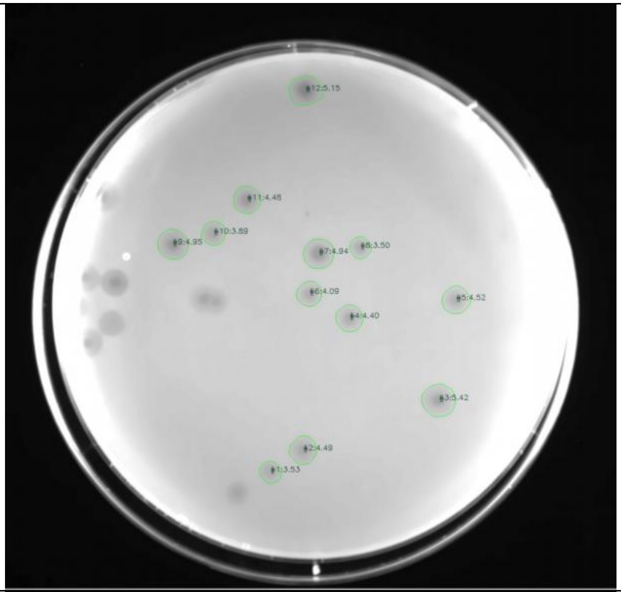 |
| **Plate_13.tif** | 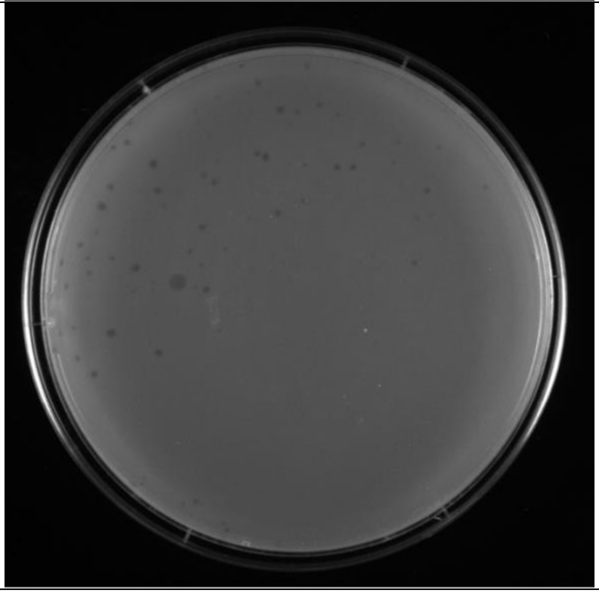 | 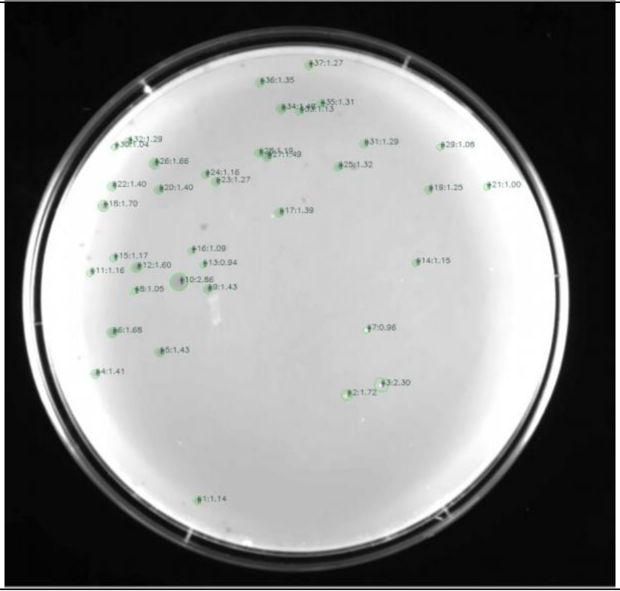 |

| | | |
|---|---|---|
| **Plate_14.tif** | 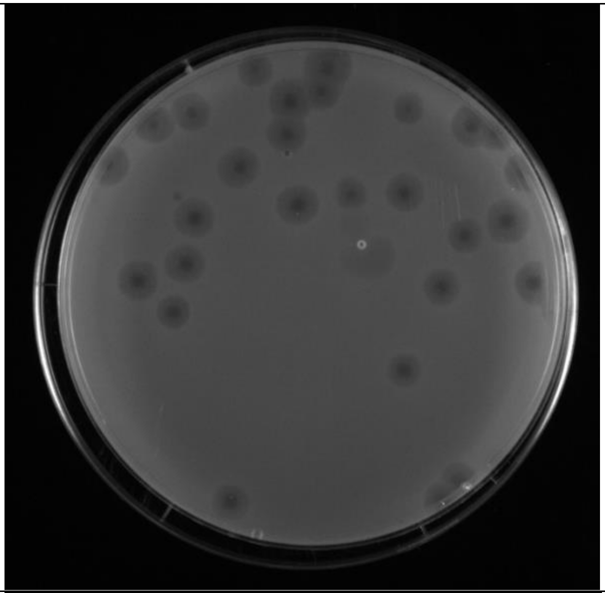 | 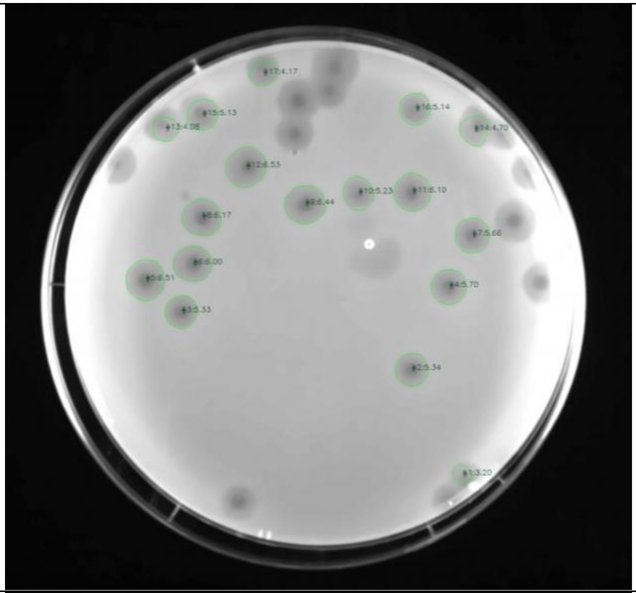 |
| **Plate_15.tif** | 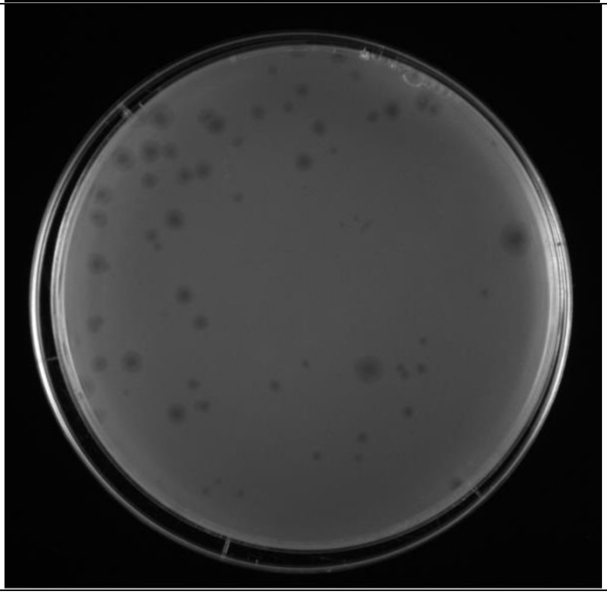 | 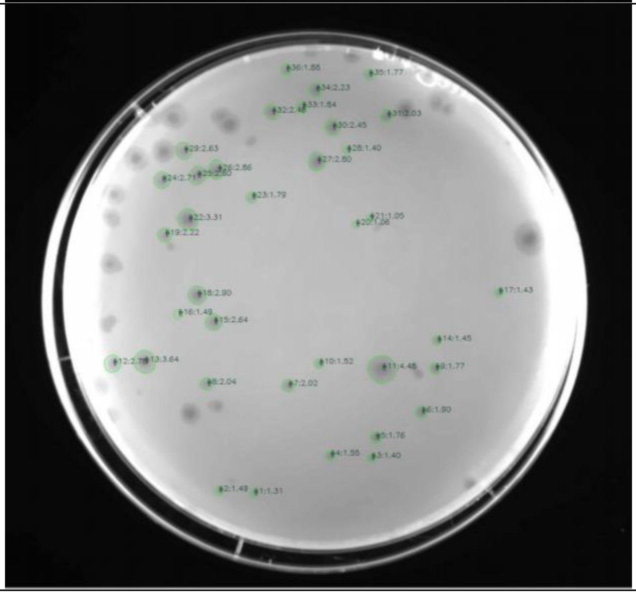 |

| | | |
|---|---|---|
| **Plate_16.tif** | 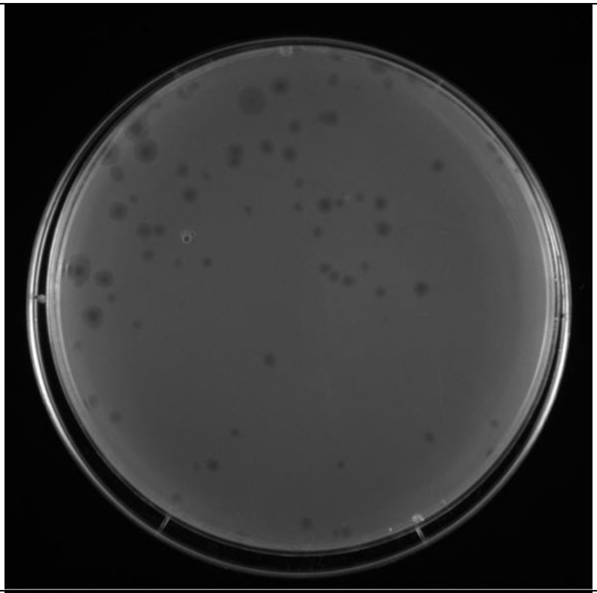 | 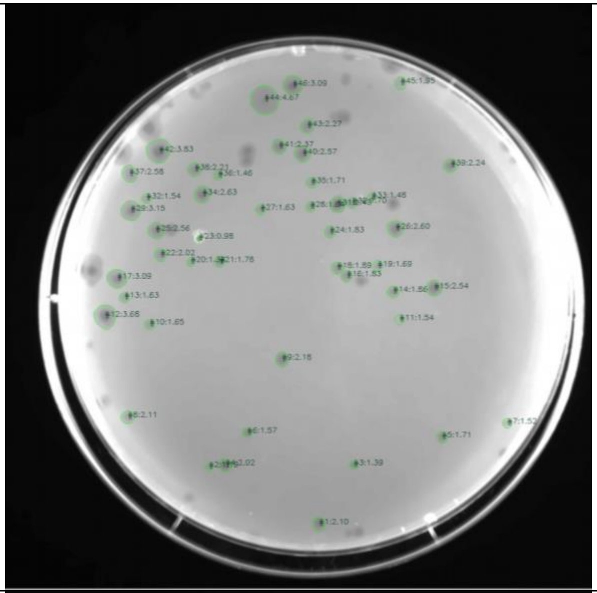 |
| **Plate_17.tif** | 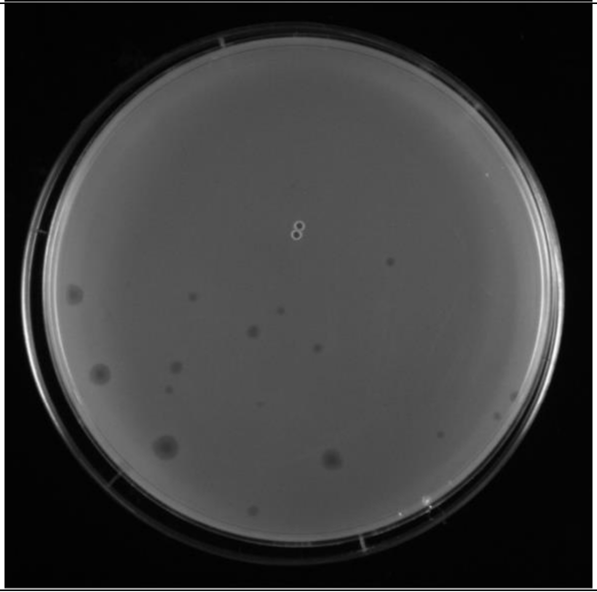 | 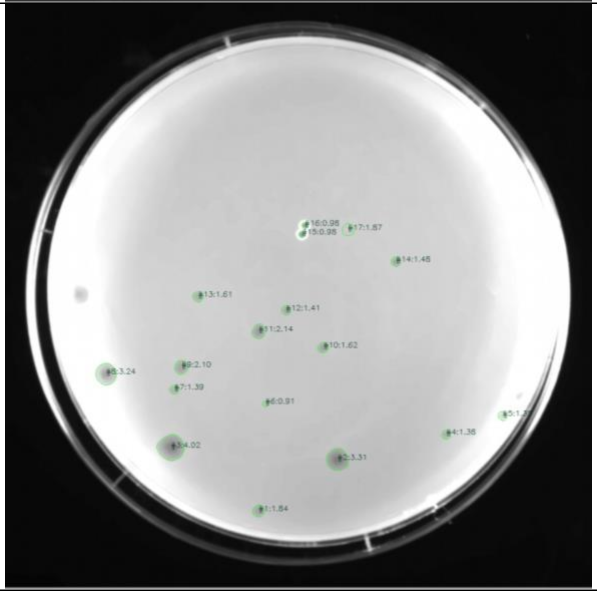 |
