## Supplementary material for "Plaque Size Tool: an automated plaque analysis tool for simplifying and standardising bacteriophage plaque morphology measurements": File S1

### Plaque Size Tool User Manual

Table of Contents

**1. INTRODUCTION ... 1**

**2. PREREQUISITES INSTALLATION ... 2**

**2.1 PYTHON INSTALLATION ... 3**
**2.2 PIP3 INSTALLATION ... 5**

**3. PLAQUE SIZE TOOL INSTALLATION ... 6**

**3.1 GITHUB ARCHIVE DOWNLOAD ... 6**
**3.2 INSTALLATION USING PIP ... 6**

**4. PLAQUE SIZE TOOL USAGE ... 7**

**4.1 PLAQUE SIZE TOOL USAGE OPTIONS ... 7**
**4.2 SINGLE FILE PROCESSING ... 9**
EXAMPLES ... 8
**4.3 BATCH FILES PROCESSING ... 10**
EXAMPLE ... 10
**4.4 OUTPUT FILES ... 11**

#### 1. Introduction

Plaque Size Tool is an open-source application written in Python 3 that is able to detect and measure bacteriophage plaques on a Petri dish image.
The source files are located at https://github.com/ellinium/plaque_size_tool.

The tool can be installed on any operation system supporting Python.
The installation guide is provided for two most frequently used OS – Windows and MacOS.

To execute installation commands on Mac use ‘Terminal’ or any other command line interpreter (CLI) preferred, on Windows use ‘Command Prompt’ (or any other CLI preferred).

The CLI screenshots taken for this manual were made on macOS High Sierra and Windows 10.

Plaque size tool was tested on the Python versions 3.7, 3.8 and 3.9.4, and if you are experiencing any problems with the higher versions, please send an email with the error to or create an issue at https://github.com/ellinium/plaque_size_tool/issues (requires registration on GitHub).

#### 2. Prerequisites installation

*Python 3.6* or higher and *pip3* should be installed on the system.

It is possible to test whether they are installed on your OS by executing a command in Terminal (macOS) or Command Prompt (Windows):

MacOS: execute ‘`python3 --version`’

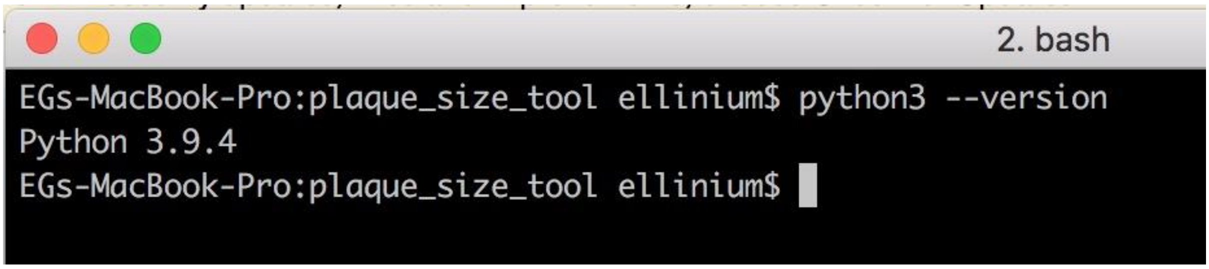


Windows: execute ‘`py --version`’

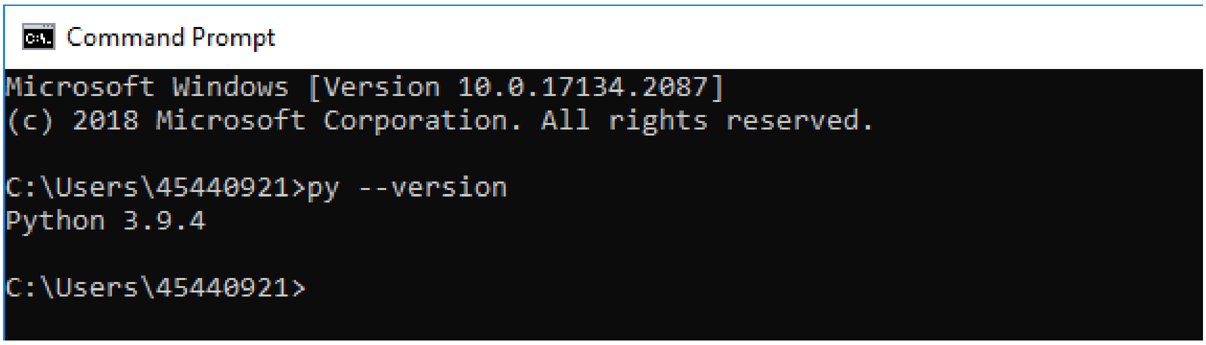


*pip3* should be installed on the system.

Pip3 is usually already installed on your system if you are using Python 3.6 or higher.
To check whether it is installed or not, execute the command ‘`pip3`’ in your CLI (the same command is used both for Windows and MacOS).

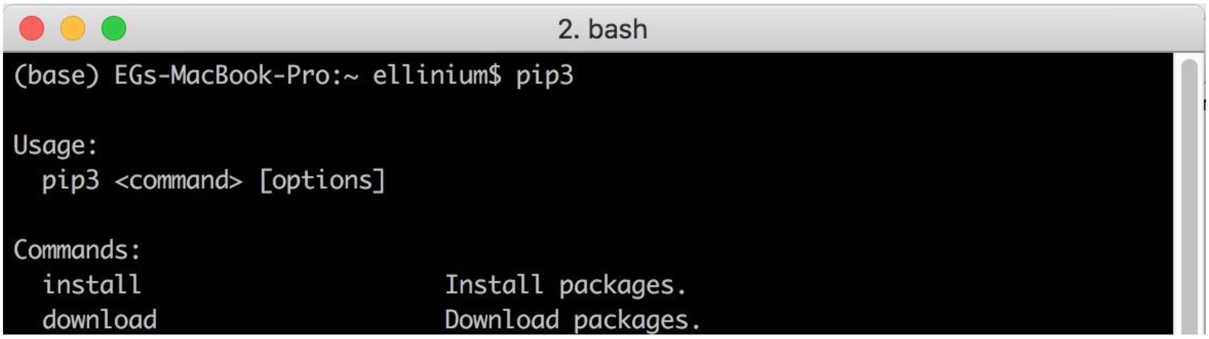


#### 2.1 Python installation

If Python3 is not found in your system, please navigate to https://www.python.org/downloads/.
On the main page there is a link to the latest version of Python3 depending on your OS.

MacOS:

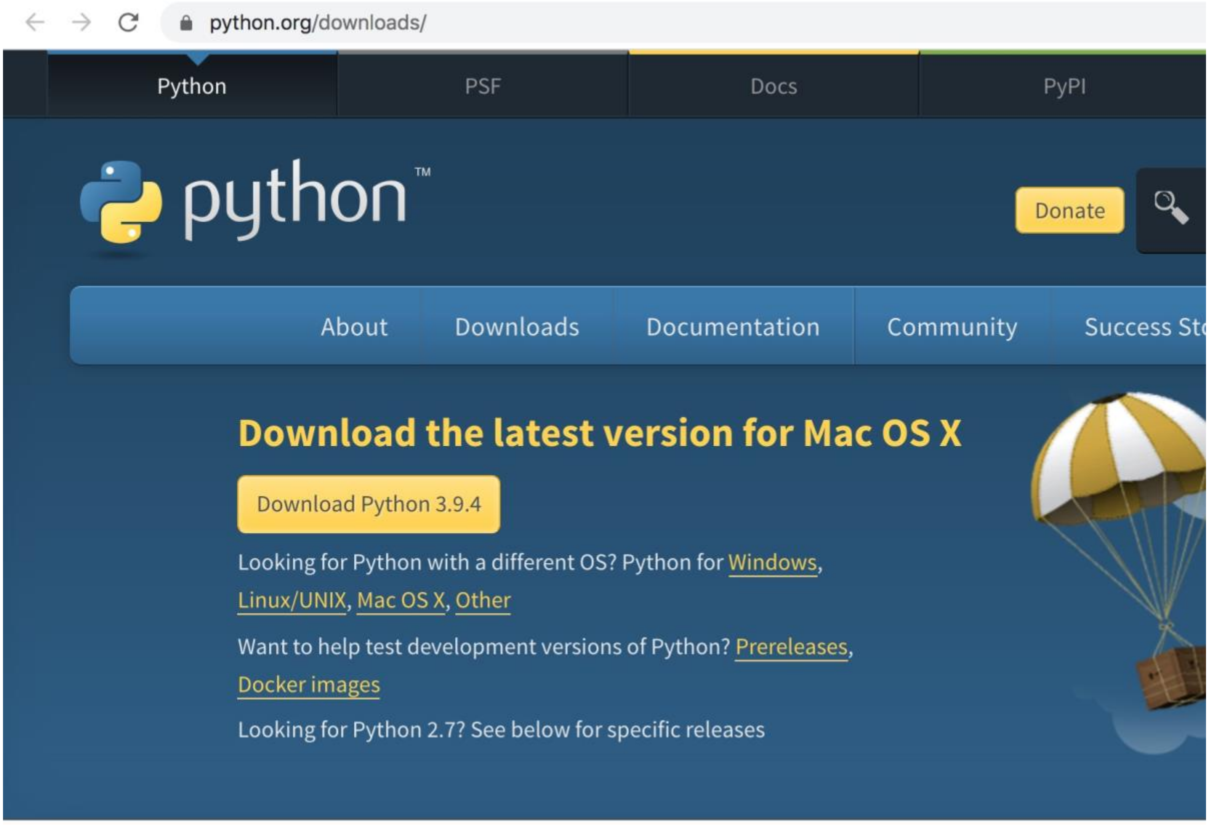

Windows:

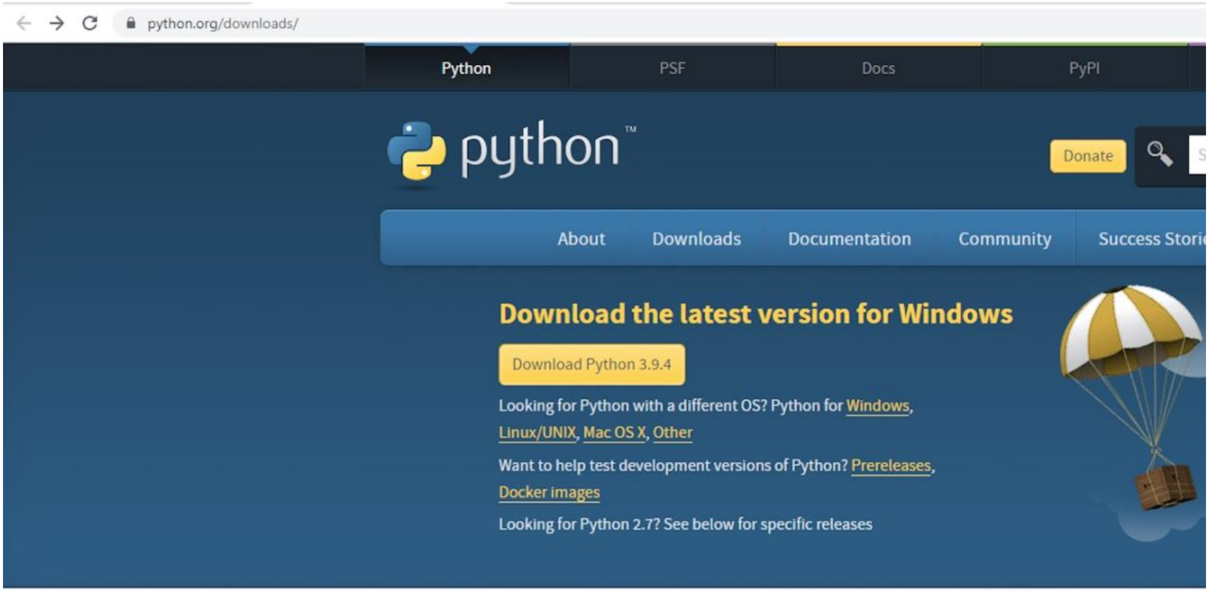


Install the latest version of Python by clicking ‘Download Python <latest version>’ and running the downloaded package.

The detailed instructions for Python download and installation are also provided at https://wiki.python.org/moin/BeginnersGuide/Download.

After Python installation, check that Python3 and pip3 were installed properly by executing the same commands as above:

MacOS:

```
python3 --version
```

```
pip3
```

Windows:

```
py --version
```

```
pip3
```

#### 2.2 Pip3 installation

If Python3 is installed in your system but *pip3* is missing, it is possible to install it separately.

For that, navigate to [https://pip.pypa.io/en/stable/installing/](https://pip.pypa.io/en/stable/installing/) and the section **'Installing with get-pip.py'**.

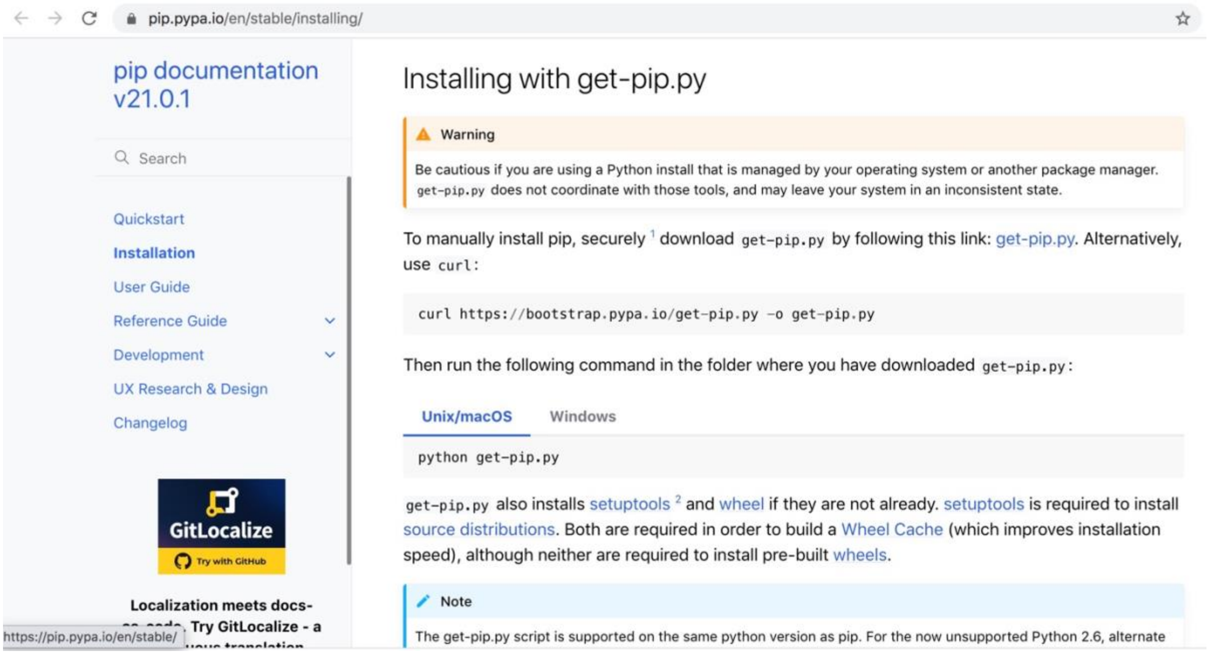


Execute the following command to download get-pip.py file:

MacOS: `python3 get-pip.py`

Windows: `py get-pip.py`

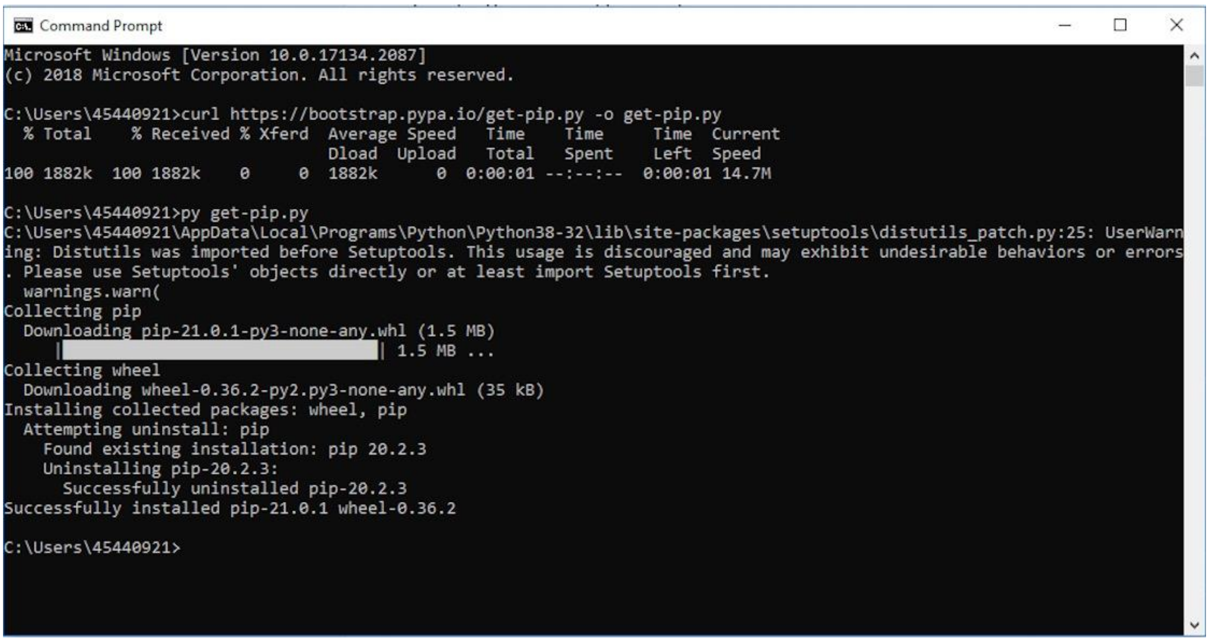


If pip3 is installed but outdated, please upgrade it by executing the following command:

MacOS: ‘`python3 -m pip install --upgrade pip`’
Windows: ‘`py -m pip install --upgrade pip`’

### 3. Plaque Size Tool installation

#### 3.1 GitHub archive download

Navigate to https://github.com/ellinium/plaque_size_tool. After that, click the green button 'Code' in the right corner and select the option 'Download Zip'.

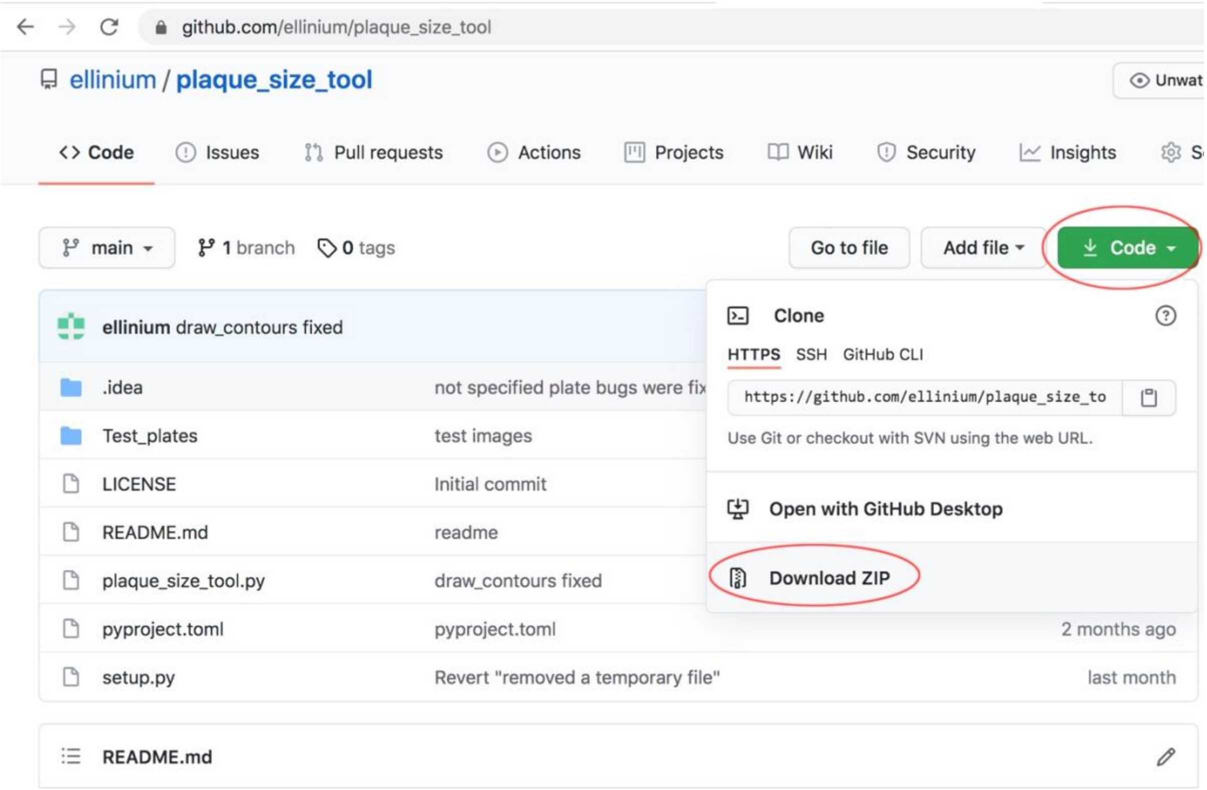


The archive called ‘plaque_size_tool-main.zip’ will be downloaded. Unpack the archive into the directory of your choice.

< OPTIONAL >: If you have already installed the program git, another option to download the files is to use the command:

`git clone https://github.com/ellinium/plaque_size_tool <directory>`, where directory is a directory for Plaque Size Tool on a local machine.

#### 3.2 Installation using pip

Next, within Terminal (macOS) or Command Prompt (Windows) navigate to the directory you unpacked the downloaded zip file into. For example, if you unpacked ‘plaque_size_tool-main.zip‘ into the ‘/home/plaque_size_tool/ plaque_size_tool-main’ directory, navigate to this directory. To confirm you are in the directory containing the plaque size tool files, type ‘ls’ on Terminal (MacOS) or ‘dir’ on Command Prompt (Windows). You should see the following files listed:

Test_plates
LICENSE
plaque_size_tool.py
pyproject.toml
README.md
setup.py

Once in the directory containing plaque size tool, execute the following command which installs all required libraries for Plaque Size Tool:

`pip3 install plaque-size-tool`

If the pip3 command worked properly you should see something like this on your screen:

```
2. bash
(base) EGs-MacBook-Pro:~ ellinium$ pip3 install plaque-size-tool
Collecting plaque-size-tool
  Using cached plaque_size_tool-1.0.7-py3-none-any.whl (11 kB)
Collecting numpy
  Using cached numpy-1.20.2-cp39-cp39-macosx_10_9_x86_64.whl (16.1 MB)
Collecting imutils
  Using cached imutils-0.5.4.tar.gz (17 kB)
Requirement already satisfied: opencv-python in /Library/Frameworks/Python.framework/Versions/3.9/lib/python3.9/site-pac
kages (from plaque-size-tool) (4.5.1.48)
Collecting pandas
  Using cached pandas-1.2.3-cp39-cp39-macosx_10_9_x86_64.whl (10.7 MB)
Collecting Pillow
  Using cached Pillow-8.2.0-cp39-cp39-macosx_10_10_x86_64.whl (2.8 MB)
Requirement already satisfied: python-dateutil>=2.7.3 in /Library/Frameworks/Python.framework/Versions/3.9/lib/python3.9
/site-packages (from pandas->plaque-size-tool) (2.8.1)
Requirement already satisfied: pytz>=2017.3 in /Library/Frameworks/Python.framework/Versions/3.9/lib/python3.9/site-pack
ages (from pandas->plaque-size-tool) (2021.1)
Requirement already satisfied: six>=1.5 in /Library/Frameworks/Python.framework/Versions/3.9/lib/python3.9/site-packages
 (from python-dateutil>=2.7.3->pandas->plaque-size-tool) (1.15.0)
Using legacy 'setup.py install' for imutils, since package 'wheel' is not installed.
Installing collected packages: numpy, Pillow, pandas, imutils, plaque-size-tool
    Running setup.py install for imutils ... done
Successfully installed Pillow-8.2.0 imutils-0.5.4 numpy-1.20.2 pandas-1.2.3 plaque-size-tool-1.0.7
(base) EGs-MacBook-Pro:~ ellinium$ 
```

MacOS and Windows use the same command for installation.

### 4. Plaque Size Tool usage

#### 4.1 Plaque Size Tool usage options

The tool can be run on a single image file (TIF, TIFF, JPG, JPEG, PNG image formats are supported) or on a directory containing several image files. The output of Plaque Size Tool will be placed into a sub-directory called ‘out’ within the /plaque_size_tool-main directory.

You can execute the command to run Plaque Size Tool from the directory used in **Installation Step 3.1.** If your current directory differs, you need to include a full path to the tool (see below for examples).

***Input parameters:***

-i to process a single image file

-d to process a directory with image files

-p is an optional parameter for the Petri dish size in millimetres

-small an optional flag, is recommended to use when the plaques are less than 2.5 mm or images are of low resolution and size

#### 4.2 Single File processing

MacOS: `python3 PATH_TO_PST/plaque_size_tool.py -i PATH_TO_THE_IMAGE_FILE [-p plate_size] [-small]`

Windows: `py PATH_TO_PST/plaque_size_tool.py -i PATH_TO_THE_IMAGE_FILE [-p plate_size] [-small]`

When the tool is executed from the directory Plaque Size Tool is installed into, the PATH_TO_PST can be omitted:

MacOS: `python3 plaque_size_tool.py -i PATH_TO_THE_IMAGE_FILE [-p plate_size] [-small]`
Windows: `py plaque_size_tool.py -i PATH_TO_THE_IMAGE_FILE [-p plate_size] [-small]`

##### Examples

To use the following examples on Windows **'python3'** is required to be replaced with **'py'**.

MacOS: `python3 plaque_size_tool.py -i Test_plates/large/Plate_4.tif`
• runs the tool on the file Plate_4.tif located in the sub-directory Test_plates/large/
• creates two files within the /out directory called: 'data-green-Plate_4.csv' and 'out_Plate_4.tif'
• all results within 'data-green-Plate_4.csv' will be shown in pixels as the plate size is not specified.

`MacOS:` `python3 plaque_size_tool.py -i Test_plates/large/Plate_4.tif -p 90`
• runs the tool on the file Plate_4.tif that has a plate size 90 mm.
• the results file 'data-green-Plate_4.csv' will contain measurements in both mm and pixels.

```
EGs-MacBook-Pro:plaque_size_tool ellinium$ cd /Users/ellinium/Documents/PST/plaque_size_tool
EGs-MacBook-Pro:plaque_size_tool ellinium$ python3 plaque_size_tool.py -i Test_plates/large/Plate_4.tif
Processing Test_plates/large/Plate_4.tif
Process completed successfully
10 plaques were found

EGs-MacBook-Pro:plaque_size_tool ellinium$ python3 plaque_size_tool.py -i Test_plates/large/Plate_4.tif -p 90
Processing Test_plates/large/Plate_4.tif
Process completed successfully
10 plaques were found
```

`MacOS:` `python3 plaque_size_tool.py -i Test_plates/small/Plate_16.tif -p 90 -small`
• runs the tool on the file Plate_16.tif that has small plaques. The results on a plate will be shown in mm.

```
2. bash
EGs-MacBook-Pro:plaque_size_tool ellinium$ python3 plaque_size_tool.py -i Test_plates/small/Plate_16.tif -p 90 -
small
Processing Test_plates/small/Plate_16.tif
Process completed successfully
46 plaques were found
```

• If executing the tool while the current working directory (which can be checked with command 'pwd' on MacOS, 'cd' on Windows) is not '/plaque_size_tool-main' then the full path to the 'plaque_size_tool.py' file AND the image file must be specified, or an error will be shown because python3 cannot find the executable file and image file.

`MacOS:` `python3 /Users/paul/plaque_size_tool-main/plaque_size_tool.py -i /Users/paul/plaque_size_tool-main/Test_plates/large/Plate_4.tif`

#### 4.3 Batch files processing

MacOS: `python3 PATH_TO_PST/plaque_size_tool.py -d PATH_TO_THE_DIRECTORY [-p plate_size] [-small]`
Windows: `py PATH_TO_PST/plaque_size_tool.py -d PATH_TO_THE_DIRECTORY [-p plate_size] [-small]`

##### Example

MacOS: `python3 plaque_size_tool.py -d Test_plates/small -p 90 -small`
Windows: `py plaque_size_tool.py -d Test_plates/small -p 90 -small`

• runs the tool on the directory Test_plates/small that contains plates with small plaques (<= 2.5 mm). The results on a plate will be shown in mm.

```
EGs-MacBook-Pro:plaque_size_tool ellinium$ python3 plaque_size_tool.py -d Test_plates/small -p 90 -small
Processing Test_plates/small/Plate_1.tif
Process completed successfully
43 plaques were found

Processing Test_plates/small/Plate_9.tif
Process completed successfully
45 plaques were found

Processing Test_plates/small/Plate_11.tif
Process completed successfully
38 plaques were found

Processing Test_plates/small/Plate_13.tif
Process completed successfully
37 plaques were found

Processing Test_plates/small/Plate_16.tif
Process completed successfully
46 plaques were found
```

#### 4.4 Output files

The tool produces two output files in the ‘out’ sub-directory that is created automatically:

**out_<file_name>** an image with identified non-overlapping plaques circled with a green line, where <file_name> is the name of the original file.

If -p (plate size) parameter is specified, the results will be shown in mm. If -p is not specified, then the results are shown in pixels.

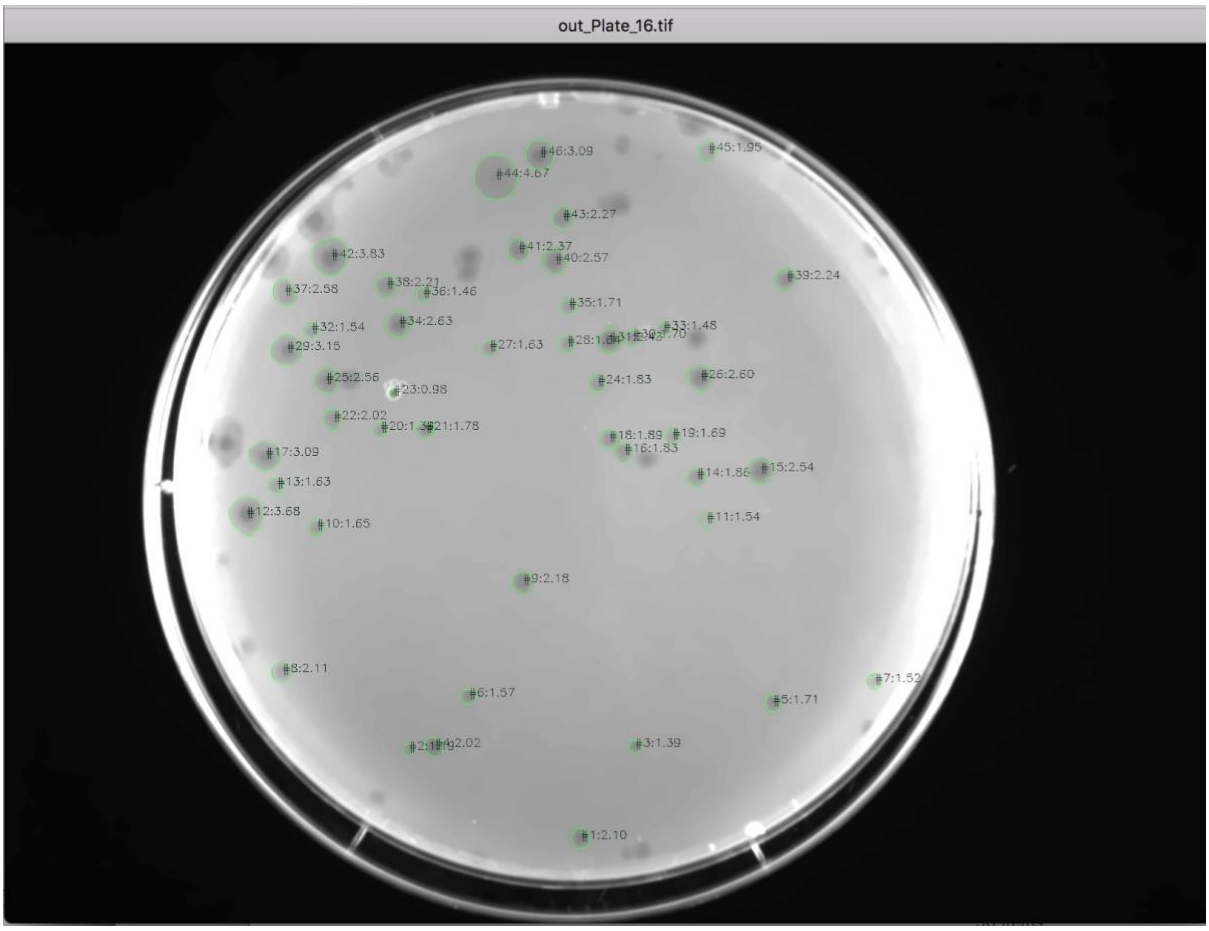

**data-green-<file_name>.csv**

a CSV file with detected plaques parameters:

INDEX_COL - the ID of the plaque that corresponds to the ID on the output image

AREA_PXL - Area of a plaque in square pixels

AREA_MM2 - Area of a plaque in square millimetres if plate size is specified

DIAMETER_PXL - Diameter of a plaque in pixels

DIAMETER_MM - Diameter of a plaque in millimetres if plate size is specified

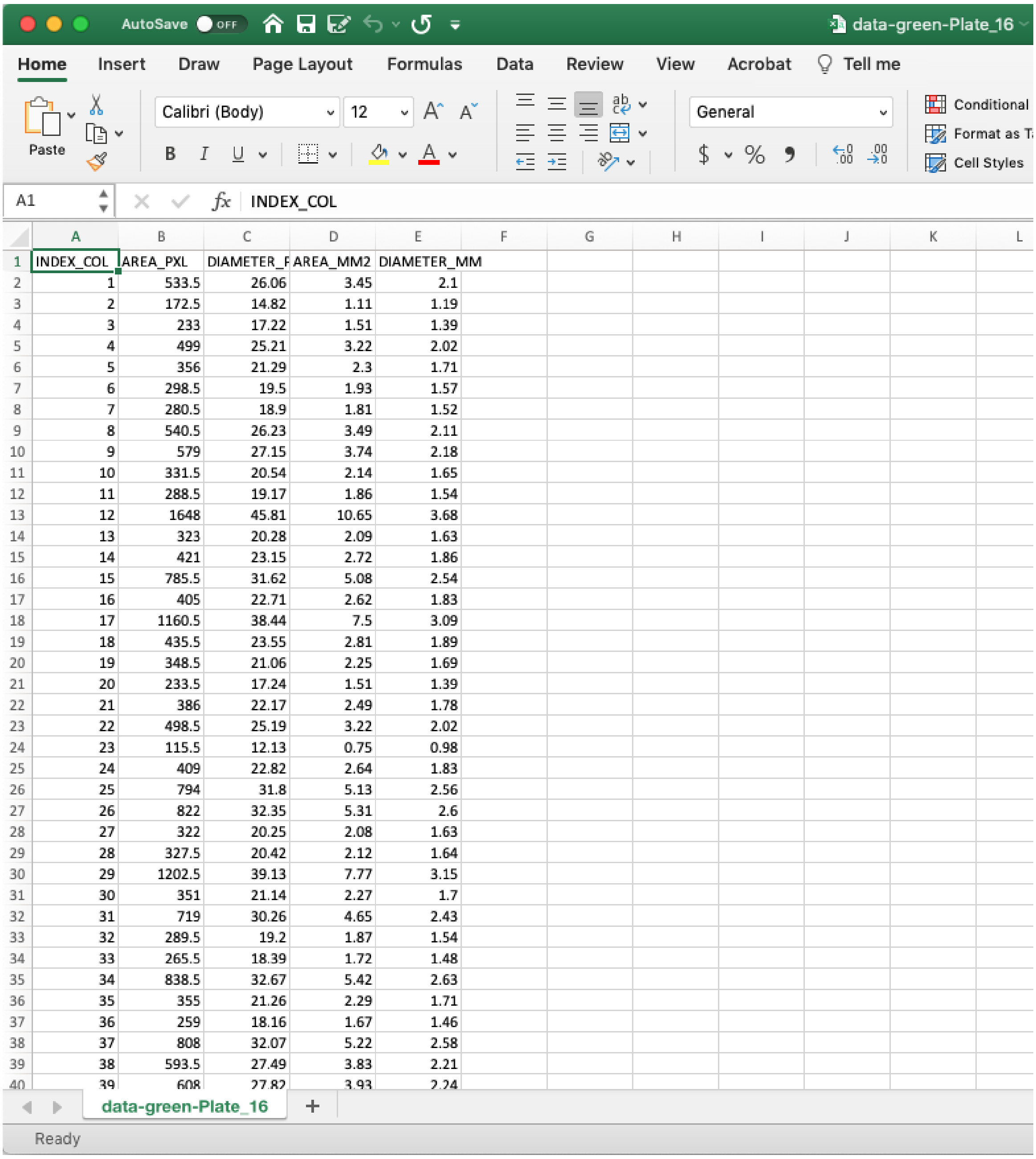


| | A | B | C | D | E |
|---|---|---|---|---|---|
| 1 | INDEX_COL | AREA_PXL | DIAMETER_F | AREA_MM2 | DIAMETER_MM |
| 2 | 1 | 533.5 | 26.06 | 3.45 | 2.1 |
| 3 | 2 | 172.5 | 14.82 | 1.11 | 1.19 |
| 4 | 3 | 233 | 17.22 | 1.51 | 1.39 |
| 5 | 4 | 499 | 25.21 | 3.22 | 2.02 |
| 6 | 5 | 356 | 21.29 | 2.3 | 1.71 |
| 7 | 6 | 298.5 | 19.5 | 1.93 | 1.57 |
| 8 | 7 | 280.5 | 18.9 | 1.81 | 1.52 |
| 9 | 8 | 540.5 | 26.23 | 3.49 | 2.11 |
| 10 | 9 | 579 | 27.15 | 3.74 | 2.18 |
| 11 | 10 | 331.5 | 20.54 | 2.14 | 1.65 |
| 12 | 11 | 288.5 | 19.17 | 1.86 | 1.54 |
| 13 | 12 | 1648 | 45.81 | 10.65 | 3.68 |
| 14 | 13 | 323 | 20.28 | 2.09 | 1.63 |
| 15 | 14 | 421 | 23.15 | 2.72 | 1.86 |
| 16 | 15 | 785.5 | 31.62 | 5.08 | 2.54 |
| 17 | 16 | 405 | 22.71 | 2.62 | 1.83 |
| 18 | 17 | 1160.5 | 38.44 | 7.5 | 3.09 |
| 19 | 18 | 435.5 | 23.55 | 2.81 | 1.89 |
| 20 | 19 | 348.5 | 21.06 | 2.25 | 1.69 |
| 21 | 20 | 233.5 | 17.24 | 1.51 | 1.39 |
| 22 | 21 | 386 | 22.17 | 2.49 | 1.78 |
| 23 | 22 | 498.5 | 25.19 | 3.22 | 2.02 |
| 24 | 23 | 115.5 | 12.13 | 0.75 | 0.98 |
| 25 | 24 | 409 | 22.82 | 2.64 | 1.83 |
| 26 | 25 | 794 | 31.8 | 5.13 | 2.56 |
| 27 | 26 | 822 | 32.35 | 5.31 | 2.6 |
| 28 | 27 | 322 | 20.25 | 2.08 | 1.63 |
| 29 | 28 | 327.5 | 20.42 | 2.12 | 1.64 |
| 30 | 29 | 1202.5 | 39.13 | 7.77 | 3.15 |
| 31 | 30 | 351 | 21.14 | 2.27 | 1.7 |
| 32 | 31 | 719 | 30.26 | 4.65 | 2.43 |
| 33 | 32 | 289.5 | 19.2 | 1.87 | 1.54 |
| 34 | 33 | 265.5 | 18.39 | 1.72 | 1.48 |
| 35 | 34 | 838.5 | 32.67 | 5.42 | 2.63 |
| 36 | 35 | 355 | 21.26 | 2.29 | 1.71 |
| 37 | 36 | 259 | 18.16 | 1.67 | 1.46 |
| 38 | 37 | 808 | 32.07 | 5.22 | 2.58 |
| 39 | 38 | 593.5 | 27.49 | 3.83 | 2.21 |
| 40 | 39 | 608 | 27.82 | 3.93 | 2.24 |
